# Architecture and self-assembly of the *Clostridium sporogenes/botulinum* spore surface illustrate a general protective strategy across spore formers

**DOI:** 10.1101/2020.01.14.906404

**Authors:** Thamarai K. Janganan, Nic Mullin, Ainhoa Dafis-Sagarmendi, Jason Brunt, Svetomir B. Tzokov, Sandra Stringer, Anne Moir, Roy Chaudhuri, Robert P. Fagan, Jamie K. Hobbs, Per A. Bullough

**Affiliations:** Krebs Institute, University of Sheffield, University of Sheffield, Sheffield S10 2TN, UK; Dept. of Molecular Biology and Biotechnology, University of Sheffield, Sheffield S10 2TN, UK; Dept. of Physics and Astronomy, University of Sheffield, Sheffield, S3 7RH, UK; Quadram Institute, Norwich Research Park, Norwich, NR4 7UA, UK; Dept. of Chemical Engineering and Biotechnology, University of Cambridge, Philippa Fawcett Drive, Cambridge CB3 0AS, UK; School of Life Sciences, Park Square, University of Bedfordshire, Luton, LU1 3JU, UK

## Abstract

Spores, the infectious agents of many Firmicutes, are remarkably resilient cell forms. Even distant relatives have similar spore architectures incorporating protective proteinaceous envelopes. We reveal in nanometer detail how the outer envelope (exosporium) in *Clostridium sporogenes* (surrogate for *C. botulinum* group I), and in other Clostridial relatives, forms a hexagonally symmetric molecular filter. A cysteine-rich protein, CsxA, when expressed in *E. coli*, self-assembles into a highly thermally stable structure identical to native exosporium. Like exosporium, CsxA arrays require harsh reducing conditions for disassembly. We conclude that *in vivo*, CsxA self-organises into a highly resilient, disulphide cross-linked array decorated with additional protein appendages enveloping the forespore. This pattern is remarkably similar in *Bacillus* spores, despite lack of protein homology. In both cases, *intracellular* disulphide formation is favoured by the high lattice symmetry. We propose that cysteine-rich proteins identified in distantly related spore formers may adopt a similar strategy for intracellular assembly of robust protective structures.

## Introduction

Spores formed by bacteria of the genera *Clostridium* and *Bacillus* provide a uniquely effective means of surviving environmental stress (1); they act as the infectious agent in pathogens such as *Bacillus anthracis, Clostridium botulinum* and *Clostridium difficile.* In the anaerobic Clostridia they are essential to survival in air. Despite the early evolutionary divergence of the genera *Clostridium* and *Bacillus*, the overall process of spore formation is strikingly similar, and a large number of the genes responsible for regulation and morphogenesis in sporulation are conserved. However, proteins making up the spore outer layers are much less conserved (2). These layers include a complex protein coat, and in some species, such as the pathogens *B. anthracis* and *C. botulinum* (but not *B. subtilis*) a distinct and deformable outermost exosporium enveloping the spore. The outer protein layers confer much of the spore’s resistance to chemical and enzymatic insult (1). The genetic control and the role of key morphogenetic proteins in *B. subtilis* spore outer layer assembly are well studied (3), but far less so in other species, particularly the Clostridia.

The exosporium defines the interface between the spore and its environment. Where the spore acts as an infectious agent, it is the first point of contact between the spore and the host. In *B. anthracis* it has roles in modulating spore germination, adhesion, protection, (reviewed in (4)), host cell uptake (5) and immune inhibition (6). The physical and structural properties of the exosporium have been best studied in the *B. cereus/anthracis* group, where it comprises a thin, continuous and hexagonally crystalline proteinaceous layer (7) (known as the basal layer) whose lattice is formed by cysteine-rich proteins ExsY and CotY (8). Its external face is decorated by a hairy nap composed of BclA, which has an internal collagen-like repeat (CLR) domain (9) that is associated with the basal layer through the ExsFA/BxpB protein (10).

In the Clostridia much less is known, with the exception of the medically important *Clostridium difficile*, where several proteins important in spore coat and exosporium assembly have been identified (11-13). Now reclassified as *Clostridioides difficile*, this species is rather distant from the main group of Clostridia however (14-16). *Clostridium botulinum* has an exosporium but its composition and assembly are poorly understood. This species is significant as a potential bioterror agent; its toxin is responsible for botulism, a severe neuroparalytic disease that affects humans, mammals and birds (17). For the highly pathogenic proteolytic strains of Group I *C. botulinum*, the closely related *Clostridium sporogenes* is a useful non-pathogenic experimental surrogate (17, 18). This makes *C. sporogenes* an attractive target for probing Clostridial spore structure and function. *C. sporogenes* exosporium is morphologically similar to that of the *B. cereus* group and has been proposed to have a hexagonally symmetric crystalline basal layer (19) and a hairy nap (20), but the detailed molecular architecture of the exosporium has not been explored. Proteins extracted from purified *C. sporogenes* exosporium (20) include, amongst others, a Clostridial-specific cysteine-rich protein, CsxA, that was detected in very high molecular weight material, together with a BclA-like protein; the latter is a possible contributor to the hairy nap, by analogy with *B. cereus*.

We now reveal the three dimensional molecular structure of a Clostridial exosporium, using electron crystallography and atomic force microscopy. The novel cysteine-rich CsxA protein has been identified as the defining structural component of the basal layer array. This provides the first detailed view of the structure of the spore envelope of *C. botulinum* Group I strains, and as CsxA homologues are encoded more widely, it will provide insights for future spore coat and exosporium research in the genus Clostridium. We show that recombinant CsxA can self-assemble into crystalline arrays identical in structure to the exosporium. Thus, we show that apparently unrelated cysteine-rich proteins from different spore-forming species can self-assemble to form remarkably similar and robust structures. We propose that diverse cysteine-rich proteins identified in the genomes of a broad range of spore formers may adopt a similar strategy of molecular tiling to build up spore structures.

## Results and Discussion

### The exosporium of C. sporogenes, and other Clostridia, is formed from a hexagonally symmetric two dimensional lattice enveloping the spore

Electron microscopy shows the exosporium enveloping the electron dense spore core; it is generally more extended at one pole (Fig. 1A). In all electron transparent areas, the exosporium appears as a thin two-dimensional crystalline layer, mostly associated with a ‘hairy nap’ and various other appendages (Fig. 1B), (20). Fourier amplitudes and phases were averaged from 5 high magnification images of negatively stained exosporium. Unit cell parameters are *a* = *b* = 110 ± 5 Å and *γ* = 120 ± 3°; phases are consistent with *p*6 symmetry. The projection map (Fig. S1A) reveals a densely stained core surrounded by a ring of 6 stain-excluding densities (black circle), separated by deeply stained pits (black rectangle). Each ring is connected to two adjacent rings by a trimeric linker (Fig. S1A; arrow). We also determined projection maps from exosporium of *C. acetobutylicum, C. tyrobutyricum, C. puniceum* and *C. pasteurianum* (Fig. S2). These all display a density distribution nearly identical to that of *C. sporogenes* (Fig. S1A).

**Figure 1.**
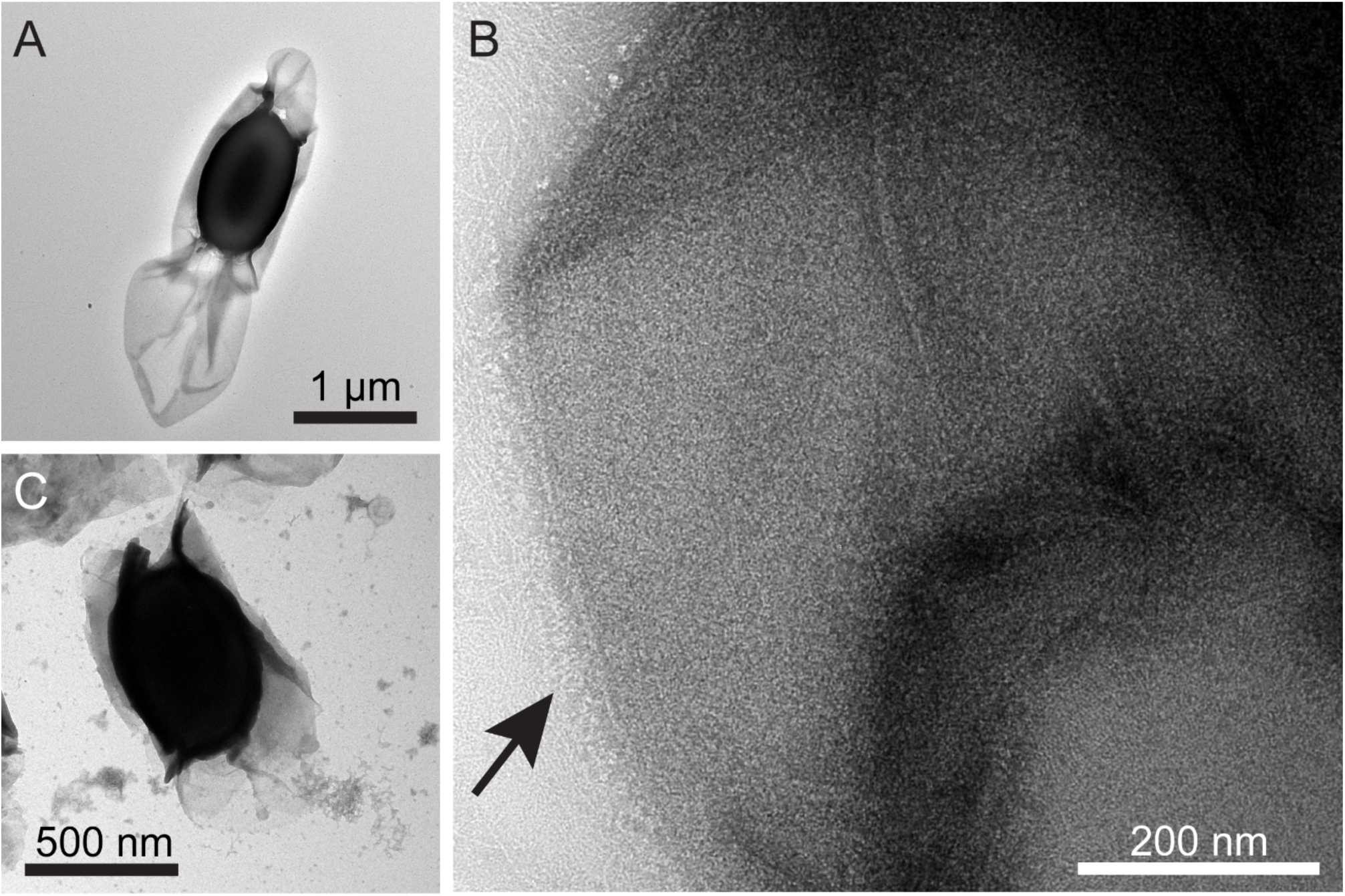
A crystalline exosporium envelops the *C. sporogenes* spore. (A) Transmission electron micrograph of a spore negatively stained with uranyl formate. The thin exosporium layer surrounds the dense spore core and is often extended at one pole. (B) A high magnification image from an area of exosporium displaying a ‘hairy nap’ fringe (arrow). (C) *csxA* mutant spore showing the core wrapped in broken sheets of material, with sloughed-off fragments in the top left corner of the image.

### Three-dimensional (3D) reconstruction of Clostridium sporogenes exosporium reveals a semi-permeable protein lattice

61 images of negatively stained exosporium from intact spores were recorded and processed in 10 separate tilt series of ±55°. 3D merging statistics are given in Table S1. In 3D (Fig. 2), the basic repeating unit is revealed as a cog-like ring with sixfold symmetry linked to adjacent rings through a small threefold symmetric bridge (arrow in Fig. 2C). The internal diameter of the ring is ∼55 Å. The outer diameter of the ring at its widest point is about ∼85 Å and the depth is ∼30 Å, similar to the thickness of the basal layer determined by AFM (see below). Major and minor pores (labelled 1 and 2 respectively in Fig. 2C) fully permeate the layer. The density revealed in this reconstruction arises only from structures that have crystalline order - any disordered features, such as surface appendages, will have been suppressed by the image averaging.

**Figure 2.**
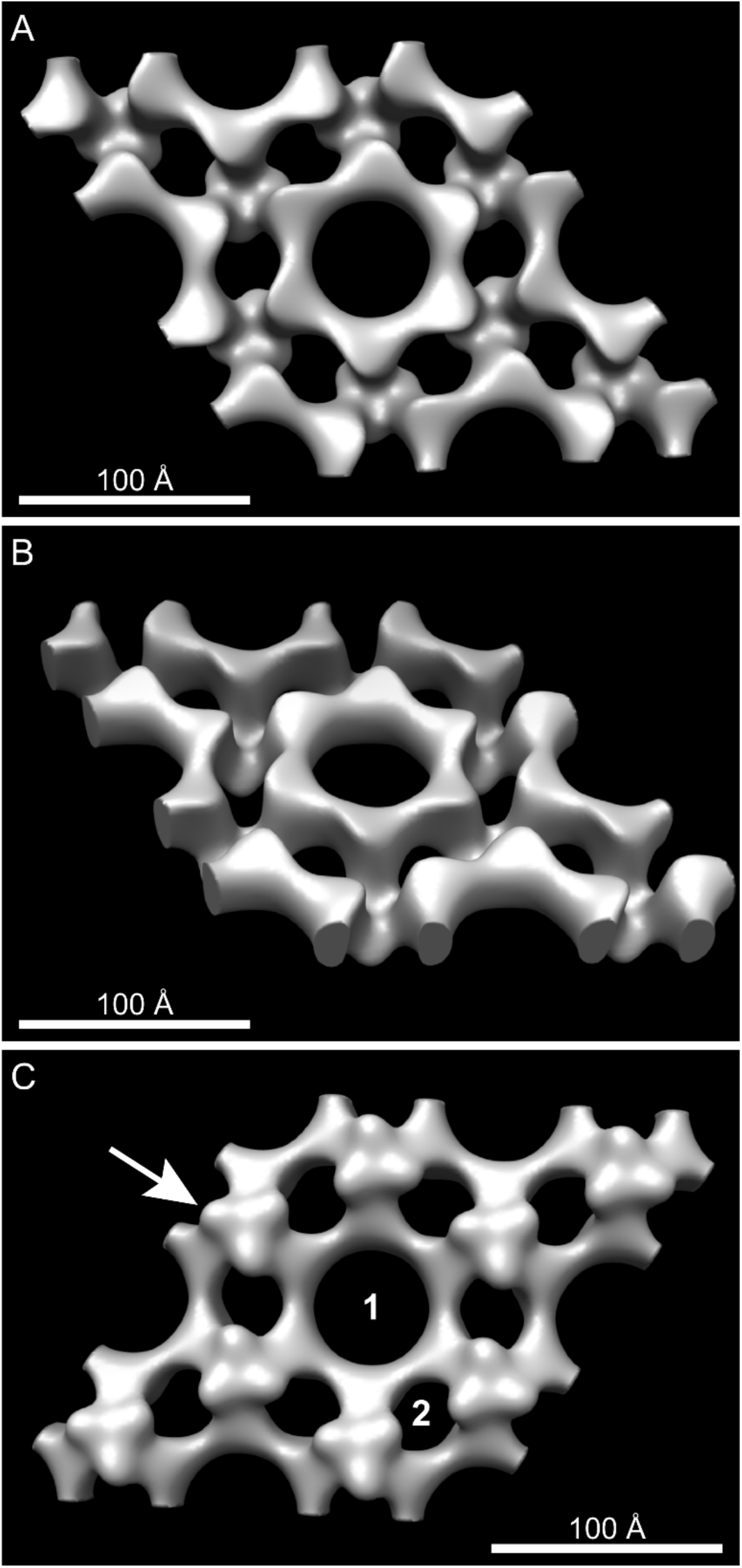
Three dimensional reconstruction of *C. sporogenes* exosporium reveals a single layered hexagonal protein lattice permeated by pores. The surface is rendered to enclose the approximate volume to match that in Fig. 4A. (A), (C) views perpendicular to the plane of the exosporium layer. (B) View at 40° to the plane. The arrow indicates the threefold symmetric linker. 1 and 2 denote central and peripheral pores. The surface shown in A corresponds to the putative exterior surface. The resolution is ∼ 25 Å.

### Atomic force microscopy (AFM) reveals opposing inner and outer faces of the exosporium

The external surface of *C. sporogenes* exosporium is largely covered by a hairy nap (Fig. 1B) and decorated with several types of appendage (20-22). As a first step in assigning the exterior face (in Fig. 2) that would bind the nap, we employed AFM to image the opposing exosporium surfaces. Exosporium fragments imaged in air displayed areas of ∼70 Å and ∼140 Å thickness (Fig. 3A, S3A), which we interpret as the thickness of single and folded over double layers of exosporium, respectively. We controlled the orientation of exosporium fragments by changing the binding conditions. When bound at pH 4, the majority of fragments presented a honeycomb-like hexagonal lattice with unit cell axes of ∼105 Å (referred to hereafter as the “honeycomb lattice”), consistent with that measured by EM (Fig. 3B). We interpret these as single layers of exosporium, bound to the substrate nap-side down, with the interior surface of the basal layer exposed to the imaging tip. Some fragments showed folded areas (2 layers thick) with a disordered surface (Fig. 3C), although occasionally a punctate hexagonal lattice with *a* = *b* = 98 ± 5 Å could be seen (Fig. 3D; white arrow). These areas are the external surface of the exosporium, with the basal layer crystal largely obscured by the hairy nap.

**Figure 3.**
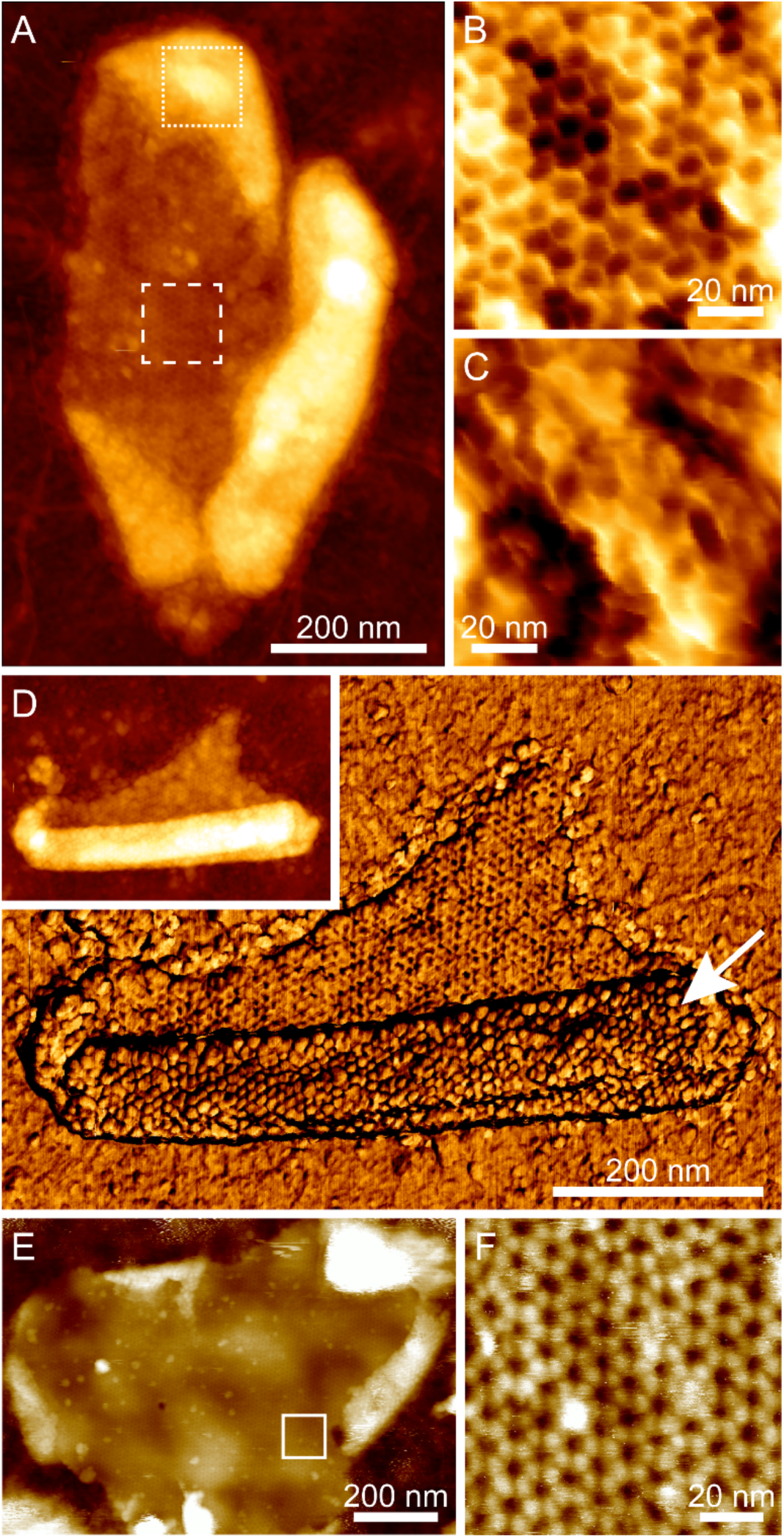
Atomic force microscopy reveals inner and outer exosporium surfaces. (A) Height image of a fragment of exosporium in air. The central area is a single layer thick (approx. 70 Å). The brighter (higher) peripheral areas are approximately twice the thickness indicating that these are folded over flaps of exosporium with the opposite surface exposed to the imaging tip. Colour scale 40 nm. (B) Higher magnification view of the area denoted by dashed box in A, showing a honeycomb-like hexagonal lattice with a unit cell of 105 ± 3 Å, proposed to be the internal surface of the basal layer. Colour scale 3 nm. (C) Higher magnification view of the area denoted by dotted box in A. This shows a disordered structure proposed to be the hairy nap on the outer surface of the exosporium. Colour scale 5 nm. (D) Inset: height image of a different exosporium fragment in air, showing a similar folded structure. Colour scale 30 nm. Main image: simultaneously acquired phase image showing the same honeycomb lattice on the single-layered part of the fragment, and a punctate hexagonal lattice with a similar unit cell size (98 ± 5 Å) on the double layered area. Colour scale 10°. (E) Height image of an exosporium fragment imaged in water. The central area is one layer thick (approx. 30 nm in water, largely due to swelling of the hairy nap). Small folded regions are visible at the edges. Colour scale 80 nm. (F) Higher magnification height image acquired in the area denoted by the box in E, showing a hexagonal lattice with a unit cell of 104 ± 3 Å, consistent with that observed in air and proposed to be the internal surface of the basal layer. Colour scale 6 nm.

When imaging in water, we again found a face with a honeycomb lattice consistent with that observed in air and and by EM, though with pronounced protrusions at the threefold symmetric vertices of each hexagon (Fig. 3E,F). The apparent thickness of exosporium in water ranged from approximately 200 to 500 Å (Fig. S3B). The greater thickness of hydrated exosporium is likely to reflect swelling of the hairy nap, though the basal layer is also thought to increase in thickness (Fig. S3C-F). Disordered regions were again observed on folded fragments, but if the imaging force was increased a punctate lattice with similar unit cell dimensions to the ordered areas in dry fragments became apparent (Fig. S4). Presumably at higher imaging forces the AFM tip was able to penetrate to the more ordered anchoring zone of the hairy nap filaments.

### The 3D structure of self-assembled recombinant CsxA matches that of the native exosporium array

The *C. sporogenes* exosporium protein CsxA is found exclusively in the very high molecular weight material extracted from *C. sporogenes* exosporium, strongly suggesting that it has a structural role in the basal layer (20). The CsxA proteins of Group I *C. botulinum* are highly similar, with amino acid identities ranging from 77% (strain A3 Loch Maree) to 100% (strain Prevot 594) (20), so data may be extrapolated more widely across the group. The *C. sporogenes csxA* gene was cloned and expressed in *E. coli*, with a C-terminal his-tag. *E. coli* cells expressing CsxA assembled stacks of sheet-like structures in the cytoplasm (Fig. S5A, white arrows). These were isolated from sonicated cells by Ni-affinity purification. The sheets were mostly folded into closed sac-like two dimensional crystals (Fig. S5B-D).

A projection map of negatively stained CsxA crystals was calculated by averaging 5 images. The unit cell parameters (*a* = *b* =111 ± 2 Å and *γ* = 120.3°± 0.4°, *p*6 symmetry (inferred from CryoEM, Table S2)) and projection structure are identical to that of native exosporium (Fig. S1B). We recorded 66 images from 11 tilt series of negatively stained 2D crystals of CsxA and calculated a 3D reconstruction as for native exosporium (shown superimposed in Fig. 4A; Table S3; Supplementary Movie 1). The structures were identical, confirming that CsxA is the core protein component of the exosporium basal layer lattice.

**Figure 4.**
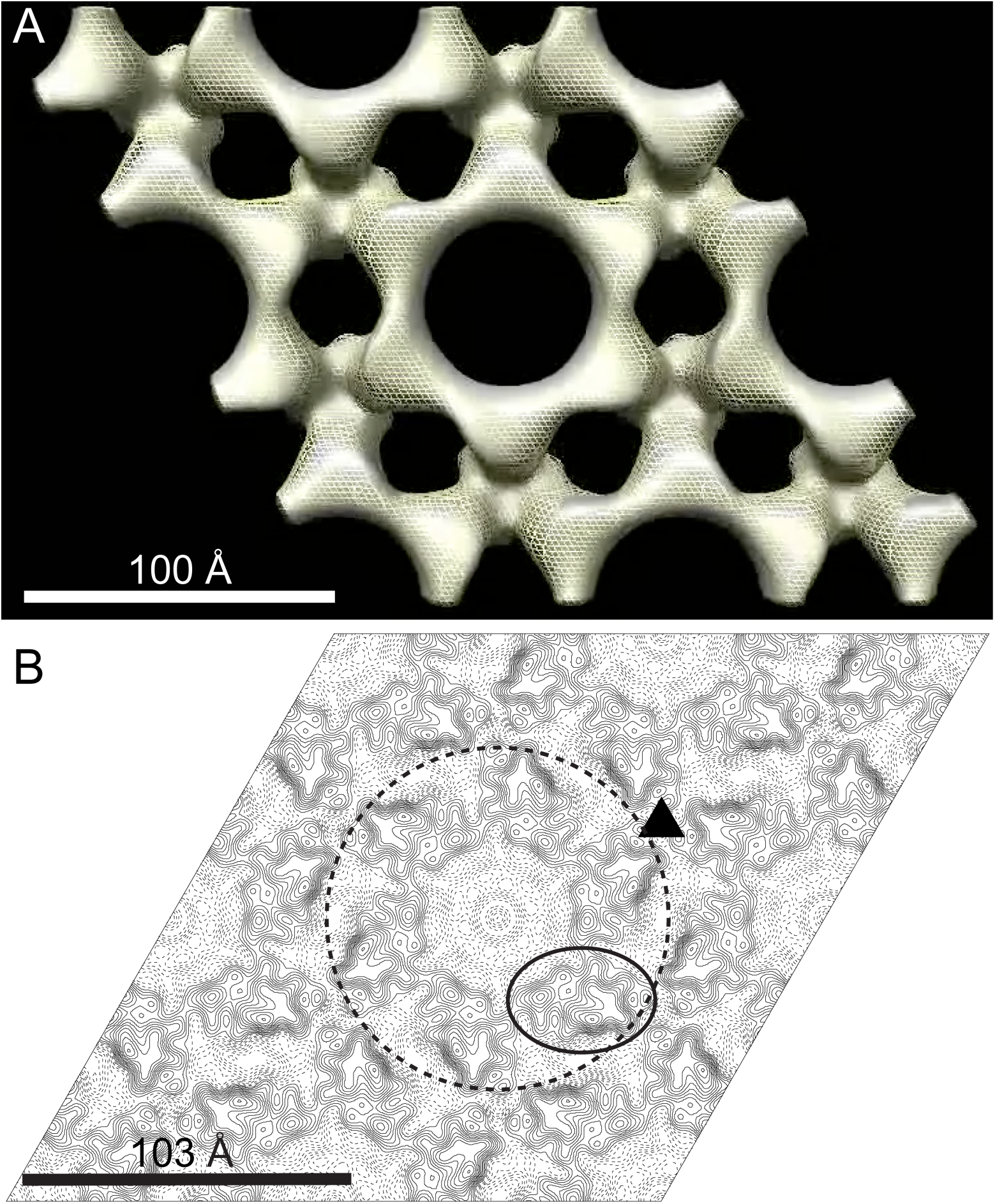
The 3D structure of the recombinant CsxA crystal is superimposable on that of native exosporium. (A) CsxA density superimposed on that of native exosporium. Surfaces are rendered at a density threshold roughly equivalent to that expected for a 33 kDa monomer. Solid grey represents native exosporium and yellow mesh represents recombinant CsxA. (B) Projection map of vitreous ice-embedded CsxA crystal at 9 Å resolution. The map is contoured at 0.2 x root-mean-square (RMS) deviation from mean density. Solid contours represent density above mean density, dashed contours represent density below the mean density. The repeating unit is a hexameric ring (dashed circle) of protein density enclosing a less dense cavity. The triangle marks the axis of threefold symmetry where rings appear connected. The density for one potential subunit is outlined.

### Electron cryomicroscopy (CryoEM) of recombinant CsxA crystals reveals a ring of six protein subunits at 9 Å resolution

The CsxA 2D crystals were better ordered than the native exosporium and more amenable to higher resolution analysis. The unit cell dimensions in vitreous ice were *a* = *b* = 103 ± 2 Å and *γ* = 120 ± 2°. Analysis of high resolution Fourier phases indicated *p*6 symmetry, thus unambiguously determining the hexameric nature of the protein assembly (Table S2). We calculated a projection map from averaged amplitudes and phases from 14 images. Phase measurements were significant to ∼9 Å resolution (Table S4). The projection map (Fig. 4B) reveals a sixfold symmetric ring of protein density with outer diameter of ∼115 Å. The ring encloses a less dense core (centre of Fig. 4B). The possible approximate envelope of one subunit is outlined and it is notable that the closest points of contact between subunits are within the hexameric ring and in the vicinity of the threefold symmetry axes that connect rings; this is consistent with the lower resolution 3D reconstruction (Fig. 2).

### AFM of CsxA crystals reveals a reversible conformational change on hydration

2D crystals of CsxA lack the external decoration present on native exosporium fragments, allowing both surfaces of the basal layer to be imaged with AFM. Two distinct surface structures, both with unit cell parameters of ∼110 Å, were observed in AFM images of hydrated CsxA crystals. One surface showed a ‘honeycomb’ lattice of pits (Fig. 5A) similar to that seen in native exosporium (Fig. 3F), meaning that we can confidently assign it to the internal surface of the basal layer. The other (external) surface displayed a lattice of hexameric assemblies with petal-like lobes (Fig. 5B). When samples were dried and imaged in air, we observed little difference in the overall arrangement of the 110 Å lattice of pits on the honeycomb face except that the threefold linkers were less pronounced, as in dehydrated native exosporium (Figs. 3B, 5C). On the ‘petal’ face, instead of a ∼110 Å lattice, we observed an array of pits with apparent ∼50 Å spacing (Fig. 5D). However, the Fourier transform showed weak first order spots indicating the true unit cell was still ∼110 Å. The overall sheet thickness of crystals decreased from ∼65 Å in water to ∼40 Å in air, regardless of which surface was exposed to the AFM tip. A cycle of dehydration followed by rehydration on crystals displaying the ‘petal’ face, showed the structural change to be reversible (Fig. S6). The mechanism and functional implications of this change remain to be determined. However, extrapolating to the native exosporium, both hydrated and dehydrated basal layer structures are likely to represent *in vivo* states, reflecting the different environments which spores would experience, such as dry to water-saturated soils or surfaces, and inside infected hosts or predators.

**Figure 5.**
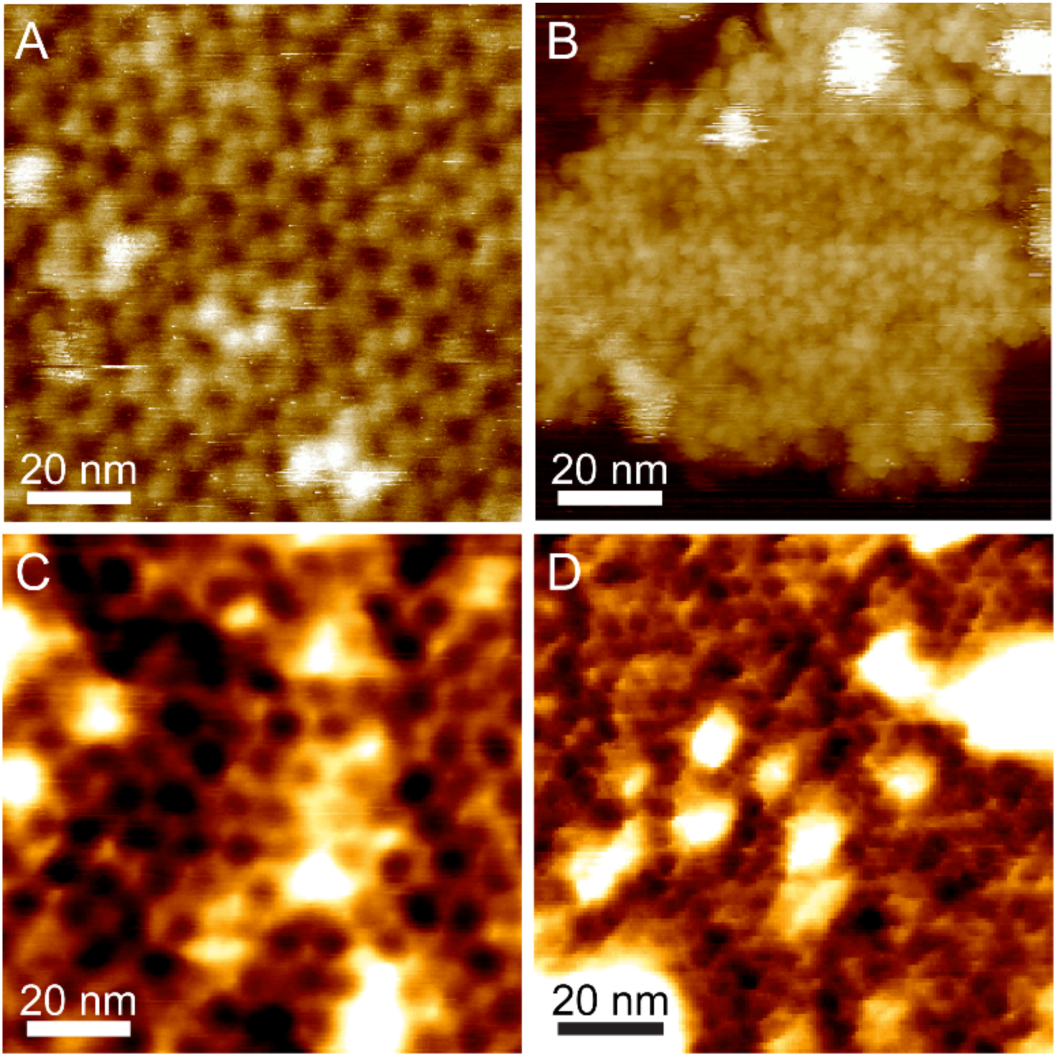
AFM of recombinant CsxA crystals reveals the external surface of the basal layer and a conformational change depending on hydration. (A) Height image of recombinant CsxA 2D crystal fragment taken in water, showing a honeycomb-like lattice with a unit cell of 103 ± 8 Å. Colour scale 6 nm. (B) Height image of recombinant CsxA 2D crystal fragment taken in water, showing a hexagonal lattice of flower like structures with a unit cell of 100 ± 6 Å. Colour scale 15 nm. (C) Height image of a recombinant CsxA 2D crystal fragment taken in air, showing a honeycomb-like lattice with a unit cell of 102 ± 2 Å. Colour scale 2.5 nm. (D) Height image of a recombinant CsxA 2D crystal fragment taken in air, showing a lattice with an apparent spacing of 50 ± 4 Å. Colour scale 2.5 nm.

### Assignment of internal and external surfaces to the 3D EM reconstruction

By comparing the EM reconstructions of native exosporium and CsxA crystals (Figs. 2 and 4A) with the respective AFM images (Figs. 3 and 5), we can tentatively assign the surface of the *C. sporogenes* EM reconstruction of native exosporium (Fig. 2) as follows: the ∼105 Å honeycomb lattice observed on the internal surface of dehydrated native exosporium fragments by AFM (Fig 3B) coincides with the face showing the raised trimeric linkers in the EM reconstruction (Fig 2C). In some cases raised features were observed by AFM at the expected linker locations (Fig. 5C). These became more prominent when imaging in water (Fig 3F, 5A). We propose that the apparent ∼50 Å lattice of pits observed on the other face of dehydrated CsxA crystals by AFM (Fig. 5D) corresponds to the larger central pores and smaller pores around the periphery of the raised hexamer surface (Figs. 2A,B, 4A). We suggest that, when hydrated, the raised hexamer swells and dominates the topography, giving rise to the “petals” lattice observed by AFM in liquid.

### CsxA arrays are thermally stable except under harsh reducing conditions

CsxA crystals were exposed to a variety of denaturing conditions (Table S5). As complete disassembly of CsxA crystals requires boiling in the presence of 2M DTT, it is likely that disulphide bonding between some or all of CsxA’s 25 cysteines plays a critical role in holding together the crystal lattice. The projection map of ice-embedded CsxA (Fig. 4B) suggests that the six monomers within a single hexameric ring are closely packed and may be connected by multiple disulphide bonds. However, the packing between rings appears less tight and it is likely that cross links occur only in the vicinity of the threefold symmetric bridge (Fig. 4B, triangle). This could explain why DTT treatment of CsxA crystals at room temperature leads to increased crystal disorder but not complete disassembly (Fig. S7).

### The CsxA protein is essential for formation of the exosporium

A *csxA* mutant of *C. sporogenes* was constructed in the more genetically amenable strain ATCC15579. Spores of this mutant no longer have a typical 110 Å lattice exosporium, consistent with the interpretation that CsxA is the core structural component of the outermost exosporium basal layer. Instead, spores appeared partially wrapped in thin broken sheets of material, fragments of which were frequently sloughed off the spores (Fig. 1C). These sheets formed 2D crystals, but with a trigonal rather than hexagonal unit cell of ∼65 Å on a side (Fig. S8). This loose proteinaceous layer may derive either from the coat or from some additional exosporium sub-layer normally more tightly associated with the spore core (23).

### Genes encoding cysteine-rich exosporium proteins and collagen-like repeat (CLR) domain proteins are colocated in distantly related spore-formers

The CsxA protein is conserved across *C. sporogenes* and related Group I *C. botulinum* species (20), and CsxA homologues are also present in a wide variety of other Clostridium species (Fig. S9). In *C. sporogenes* strain ATCC 15579, and others, the CsxA protein is encoded in a gene cluster between *zapA* and a gene encoding a U32 family peptidase (Fig S10). In ATCC 15579, the cluster also encodes a glycosyl transferase and proteins containing collagen-like repeats (CLR domains), including the BclA protein. BclA is found associated with CsxA in high molecular weight exosporium protein complexes, and the CLR domain protein, BclB, is detected in bulk exosporium (20). By analogy with *B. cereus* and *B. anthracis*, at least some of these CLR domain proteins are likely glycoproteins forming the hairy nap on the surface of the exosporium. This is reminiscent of *B. anthracis*, in that genes for the three proteins that are found in the high molecular weight exosporium complexes (BclA, ExsFA and the cysteine-rich protein, ExsY) are encoded in a cluster, along with those encoding glycosyl transferase and other genes (24). In *C. sporogenes*, no other identified exosporium proteins are encoded at this position, although other conserved ORFs may be present.

Gene synteny, but with variations in the CLR-encoding ORFs, is found in other *C. sporogenes* and Group I *C. botulinum* strains (Fig. S10); more distant *csxA* homologues (e.g. in *C. tyrobutyricum, C. puniceum*, and *C. beijerinckii*) are also frequently colocated with genes encoding CLR proteins and glycosyl transferases, though not necessarily in the same genomic context. More generally, CLR domain proteins and cysteine-rich proteins can be found encoded within 20 kb of each other across many spore formers (Supplementary data set 1); a number of these cysteine-rich proteins are annotated as spore coat proteins and others are potential candidates for spore coat and/or exosporium proteins.

### Distantly related spore formers use different proteins but adopt similar design principles to build the spore envelope

The representatives of the Bacillales and Clostridiales that we have studied (*B. cereus/anthracis, C. sporogenes/botulinum, C. acetobutylicum, C. tyrobutyricum, C. puniceum* and *C. pasteurianum*), possess exosporia of remarkably similar morphology, with a crystalline basal layer enveloping the spore core and decorated by a more disordered ‘hairy nap’ (Fig. 1A,B) (25); in the cases of *B. cereus* and *C. sporogenes*, a disordered outer surface and ordered inner surface are observed by AFM (Fig. 3) (7, 20). In *B. cereus*/*anthracis, C. botulinum*/*sporogenes* and other Clostridia, the basal layer of the exosporium has a regular tiling pattern of interlinked sixfold symmetric oligomers (compare Fig. 2, 3, S1 and S2 with (7)). In *B. cereus* the core components are ExsY and CotY (8); these are cysteine-rich and analogous to, but not homologous to CsxA. The lattice spacings in the Clostridia tested were higher (∼110 to ∼127 Å versus ∼80 Å in *B. cereus/anthracis*), and the *C. sporogenes* basal layer reveals an apparently more permeable structure, with pores of ∼55 Å, compared to ∼20 Å (25) (Fig. S2). Although the effective diameters for diffusion would be smaller than those physically measured in the reconstructions (26) all could allow the permeation of small molecule germinants (27, 28), but might still exclude hostile enzymes and antibodies.

The similar design of hexagonally tiled meshes across the basal layers of spore formers reflects apparently common functions of acting as semi-permeable molecular filters and in forming a platform onto which other proteins may bind, including proteinaceous appendages. In the *B. cereus* group, the CLR-domain protein BclA is attached to ExsY via an anchor protein, ExsFA/BxpB (10). There is no ExsF homologue in Clostridia, and there is no evidence of a third protein in the CsxA-BclA complexes extracted from *C. sporogenes* (20), so whether BclA is directly anchored to the CsxA basal layer is not known. Notably, a second *C. sporogenes* cysteine-rich protein, CsxB, was detected in association with BclA, but not with CsxA, in somewhat smaller complexes by SDS-PAGE, and CsxB was much more easily dissociated into likely oligomers and monomers; its role in the exosporium, like that of BclA of *C. sporogenes*, and other exosporium proteins identified by Janganan *et al.* (20), remains to be directly tested.

### Self assembling cysteine-rich proteins form robust protective layers in spores from distantly related species

Cysteine-rich proteins are emerging as a characteristic feature of the proteomes of spores (8, 13, 20, 29-32). We have now identified a variety of spore proteins, some with very different sequences, that have the common properties of being cysteine-rich and capable of self-organization into extended 2D ordered arrays resembling natural assemblies found in the native spore. It is notable that the identical crystal packing symmetry that we see in CsxA assemblies has been found in the unrelated cysteine-rich proteins, CotY from *B. subtilis* spore coat (31), and ExsY and CotY from *B. cereus* exosporium (8). Arrays of these proteins isolated from an *E. coli* expression host also require harsh reducing conditions and boiling for complete disassembly. This robust cross-linking of arrays is likely to reflect the situation in the native spore - harsh denaturing and reducing conditions are required for complete disintegration of the hexagonal lattice of *C. botulinum* and *B. cereus* exosporium (8, 33). Whilst the cellular milieu of the native mother cell, where the exosporium is assembled, or of the heterologous *E. coli* host is generally considered ‘reducing’ and intracellular disulphide bonding is rare, the ordered lattice and high symmetry of the respective proteins could provide sufficient avidity to overcome this and still drive cooperative intracellular disulphide cross-linking (Fig. 6).

**Figure 6.**
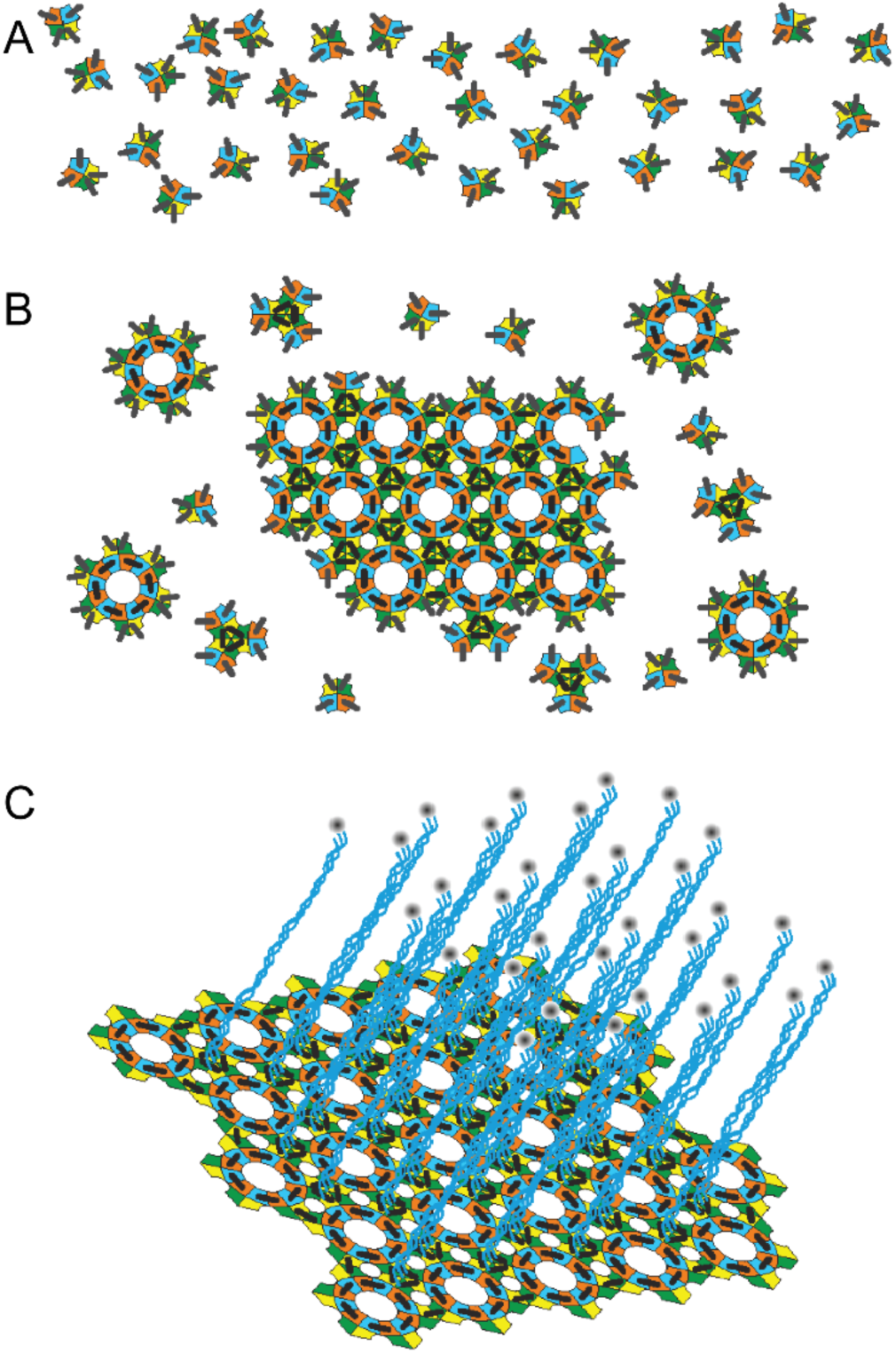
Two groups of distantly related spore formers show the same hierarchy of assembly states on the spore surface. The schematic diagram indicates the hierarchy of assembly states of exosporium in the *B. cereus* group and the *C. botulinum* group. (A) Monomeric units of ExsY in the *B. cereus* group and CsxA in the *C. botulinum* group self-assemble into (B) a hexameric disulphide cross-linked array. The high symmetry of the growing lattice would enhance the avidity for disulphide bond formation-represented by the black bars. Note that the number of cysteine residues participating in cross-linking is unknown. The colours represent schematically different surface regions of the asymmetric monomers. (C) In both groups, the surface lattice is embellished with a ‘hairy nap’, known to be mostly the CLR protein, BclA, in the case of the *B. cereus* group. In the case of the *C. botulinum* group it is likely to be the respective BclA equivalent.

As we have done previously for ExsY (8), for CsxA we can describe a hierarchy of units of assembly in the protein lattice, from monomer through hexamer to the extended cross-linked 2D hexagonal array. In both cases this array of cysteine rich proteins can then be decorated by filamentous appendages including proteins with CLR domains (Fig. 6). We propose that the many cysteine-rich coat and exosporium proteins identified in the genomes of spore formers are likely to have similar structural roles to those of CsxA, ExsY and CotY. Indeed, disulphide bonding plays a central role in the development of fully assembled spores. Reduction of spore disulphide bonds sensitizes *Bacillus* and *Clostridium* spores to lysozyme and hydrogen peroxide, thus emphasizing their protective role (34). Moreover, Aronson and Fitz-James (35) found evidence for the role of self-organisation by partially reconstituting outer coat layers of *B. cereus* that had been treated initially with DTT and urea.

It may not be a general requirement for self-assembly of cysteine-rich proteins that they form the extensive crystalline arrays that we have described; some may be only locally ordered. Nevertheless, extensive crystalline layers of unidentified proteins are associated with spore coats and exosporium-like layers in a wide range of spore forming species (36-44).

### Role of the exosporium in outgrowth

We have shown that those parts of the *in situ* exosporium that are accessible for diffraction analysis adopt the basal layer structure identical to that of recombinant CsxA crystals (Figs. 1A and 4A). However, it is possible that the spore is not completely enclosed by a uniform and continuous CsxA lattice. Although low resolution scanning electron micrographs suggest a mostly uniform exosporium fully enveloping the *C. sporogenes* spore there does appear to be a relatively weak structure (sporiduct) at one pole through which the outgrowing cell emerges (45). This may be made up of different proteins and could incorporate a ‘cap’ as seen in *B. anthracis* (46, 47). The exosporium does not appear to tear beyond this region, indicating that the CsxA lattice is relatively resistant to any pressure from the emerging vegetative cell.

### Conclusion

We have achieved the first three dimensional molecular reconstruction of a Clostridial spore surface. The *C. sporogenes* spore is enveloped by an exosporium built with a paracrystalline basal layer, the core component of which is the cysteine-rich protein, CsxA. This is the first example of a Clostridial protein capable of self assembly to form a paracrystalline cross-linked spore layer; a phenomenon that we previously demonstrated in unrelated spore surface proteins of the Bacillales (8, 31). We have demonstrated how apparently unrelated proteins from different species can assemble to form similar highly symmetric tiled arrays. The proteins are cysteine-rich and self-assemble to form highly resilient spore layers. We propose that diverse cysteine-rich proteins identified in the genomes of a broad range of spore formers may adopt a similar strategy for assembly.

## Materials and methods

A full account of the methods used is in Supplementary Information.

### Spore and exosporium preparation

For *C. sporogenes* NCIMB 701792 (NCDO1792) and all other species, spore and exosporium preparations were performed as described in (20) except that spores of *C. acetobutylicum* NCIB 8052, *C. tyrobutyricum* NCDO 1756, *C. puniceum* BL 70/20 and *C. pasteurianum* NCDO 1845 were harvested from potato extract agar.

### csxA mutant construction

A *csxA* mutant of *C. sporogenes* strain ATCC15579 was generated using the Clostron system as previously described (27). *C. sporogenes* strain NCIMB 701792 (NCDO1792) was not used for mutational studies as this strain is erythromycin resistant and not amenable to Clostron mutation.

### Expression of csxA in E. coli

The *csxA* gene was amplified by PCR from genomic DNA of NCIMB 701792, and cloned into pET21a, forming a C-terminal His_6_ fusion. After 3h of induction in BL21(DE3) pLysS, cells were harvested, sonicated and the CsxA 2D crystals recovered by binding to Ni-NTA agarose beads, washing with buffer, and elution with 1M imidazole.

CsxA crystals were examined by negative stain EM (see below) after incubation under a variety of combined conditions including 95 °C heat treatment, 8M urea and 2M DTT-see Table S5.

### Electron microscopy of E. coli cell sections

*E. coli* cells over expressing CsxA were fixed in glutaraldehyde and osmium tetroxide and embedded in propylene oxide/araldite resin. Ultrathin sections, approximately 70-90 nm thick, were stained with uranyl acetate and Reynold’s Lead Citrate. The sections were examined on a FEI Tecnai 120 G2 Biotwin microscope at 80 kV with a Gatan Orius SC 1000B camera.

### Electron microscopy of exosporium and crystals

Spores and CsxA crystals were stained with uranyl formate and examined in a Philips CM100 microscope at 100 kV. Micrographs were recorded under low dose conditions on a Gatan MultiScan 794 1k x 1k CCD camera at specimen tilts over a range of ± 55° in 10° steps.

For cryomicroscopy CsxA crystals were frozen and imaged at 200 kV in a Philips CM200 FEG TEM or a FEI Tecnai Arctica. Micrographs were collected on a Gatan UltraScan 890 2k x 2k camera or a FEI Falcon III camera respectively.

### Image processing

Electron micrographs were processed within the *2dx* software suite (48) with *p6* symmetry enforced. For data from tilted, negatively stained samples, the variation of amplitude and phase along 0,0,l was estimated from a plot of the maximum contrast on each Z-section in the 3D map. 3D surface representations were rendered with CHIMERA (49).

For frozen hydrated CsxA crystals we merged and averaged 14 images in *2dx* (48) to 9 Å resolution. A negative temperature factor (B-factor) was estimated and applied to the projection map by scaling averaged image amplitudes against bacteriorhodopsin electron diffraction amplitudes (50).

### Atomic Force Microscopy

Exosporium fragments were bound to poly-d-lysine coated cover slides at pH 4. 2D crystals of CsxA were bound to freshly cleaved mica. Samples were either imaged in water or washed, dried and imaged in air. Imaging in air was performed using a JPK NanoWizard Ultra AFM in AC mode with TESPA V2 cantilevers in a home built vibration and acoustic isolation system. Imaging in water was performed using a Bruker Dimension FastScan AFM in Tapping Mode with FastScan D cantilevers. Images were processed and analyzed using JPK DP software, Gwyydion and NanoScope Analysis.

## Supporting information

Supplementary data sets 1

## Acknowledgements

We thank Chris Hill for help with EM of thin sections. All electron microscopy work was carried out in the University of Sheffield’s Faculty of Science Electron Microscopy Facility. We also thank Paul Kemp-Russell and Simon Dixon who fabricated the acoustic enclosure for AFM. JKH and NM gratefully acknowledge the Imagine: Imaging Life initiative at the University of Sheffield and the EPSRC for financial support through its Programme Grant scheme (Grant No. EP/I012060/1). PAB and TKJ gratefully acknowledge the Wellcome Trust for financial support. ADS was in receipt of a White Rose BBSRC DTP studentship. Author contributions: T.K.J., N.M. and P.A.B. designed research; T.K.J., N.M., A. D.-S., J.B., S.B.T., S.S., R.C. and P.A.B. performed research; T.K.J., N.M., A.D.-S., A.M., R.C., R.P.F., J.K.H. and P.A.B. analyzed data; T.K.J., N.M., A.M., R.P.F., J.K.H. and P.A.B. wrote the paper.

The authors declare no competing interests.

