## Supplementary information for "Architecture and self-assembly of the *Clostridium sporogenes/botulinum* spore surface illustrate a general protective strategy across spore formers"

***Supplementary movie 1***

Comparison of EM reconstructions of native *C. sporogenes* exosporium and CsxA crystals.

***Supplementary Figures***

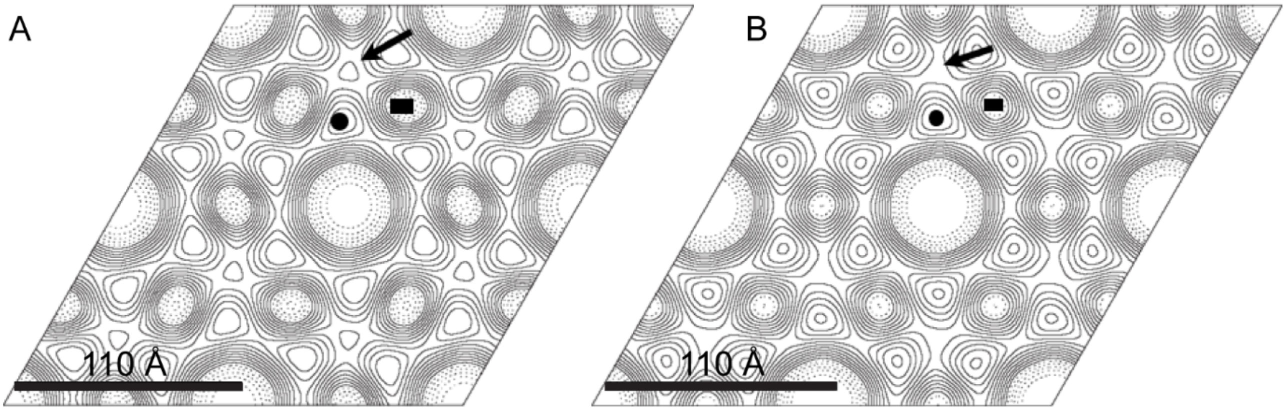


**Figure S1. Projection maps of *C. sporogenes* exosporium and recombinant CsxA crystals display hexagonal symmetry.** (A) Projection map of negatively stained *C. sporogenes* exosporium determined to ~20 Å resolution. Solid contours represent areas of stain-excluding density, corresponding to areas of higher protein density. (B) Projection map of negatively stained CsxA crystal determined to ~20 Å resolution. Black circle indicates stain-excluding density, rectangle indicates peripheral pore and arrow indicates threefold linker.

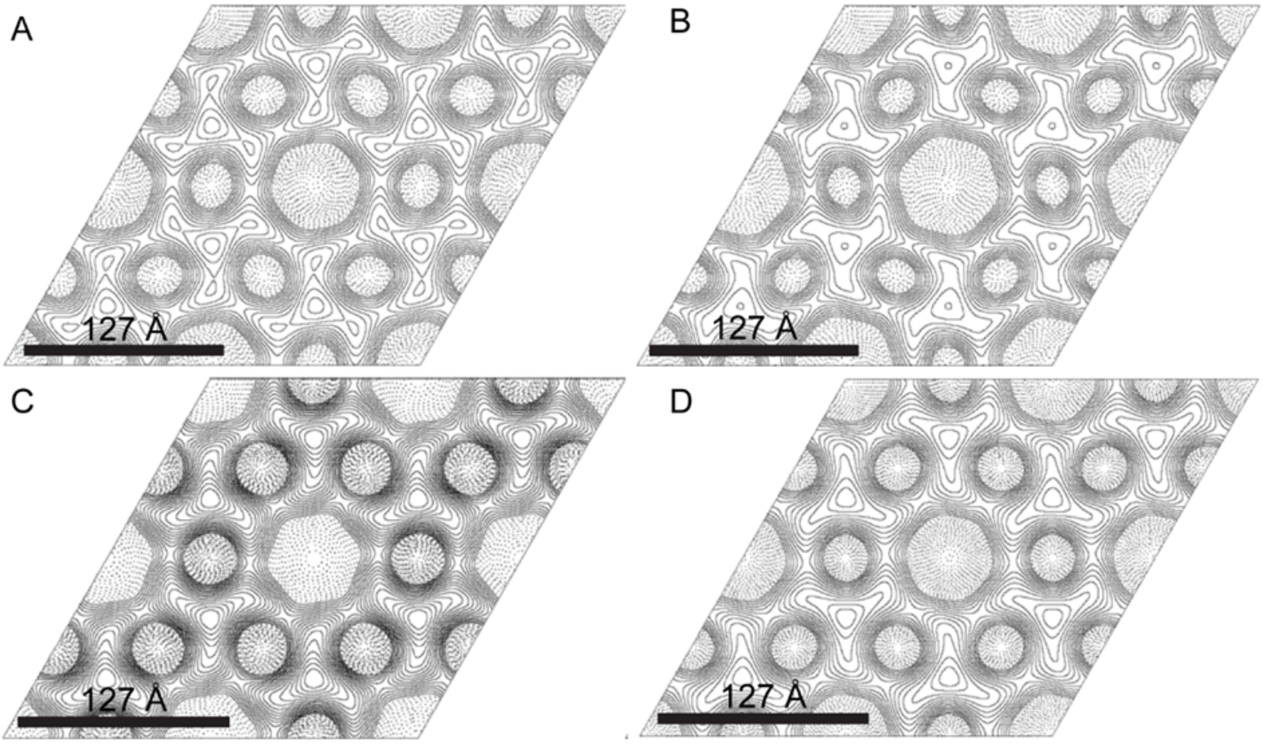


**Figure S2. Projection maps of exosporium from four *Clostridium* species show a structure similar to that of *C. sporogenes.*** See Fig. S1 legend for details. (A) *C. acetobutylicum* NCIB 8052. (B) *C. tyrobutyricum* NCDO 1756. (C) *C. puniceum.* (D) *C. pasteurianum* NCDO 1845.

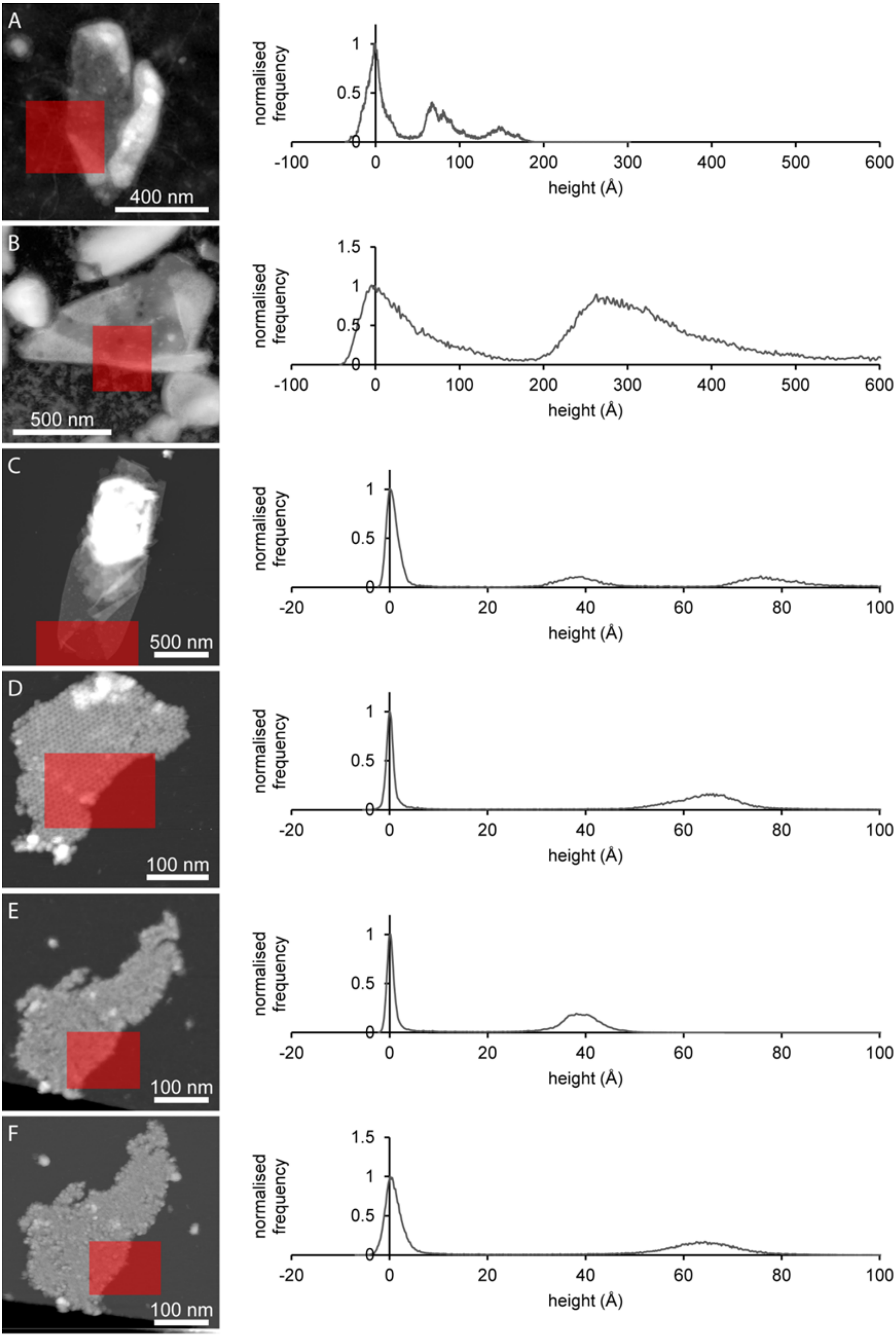
A
400 nm
normalised frequency
height (Å)
B
500 nm
C
500 nm
D
100 nm
E
100 nm
F
100 nm

**(Previous page): Figure S3. AFM thickness measurements of exosporium and CsxA crystals in air and liquid.** Left column: height images with red shading denoting areas selected for analysis. Right column: height histograms calculated for the red shaded areas. The histograms have been shifted along the height axis such that the modal value of the peak corresponding to the background substrate is at zero height. (A) native exosporium in air (greyscale 32.5 nm); (B) native exosporium in water (greyscale 177 nm); (C) CsxA crystal (internal side upwards) in air (greyscale 60 nm); (D) CsxA crystal (internal side upwards) in water (greyscale 15.6 nm); (E) CsxA crystal (external side upwards) in air (greyscale 11 nm); (F) CsxA crystal (external side upwards) in water (greyscale 21.8 nm).

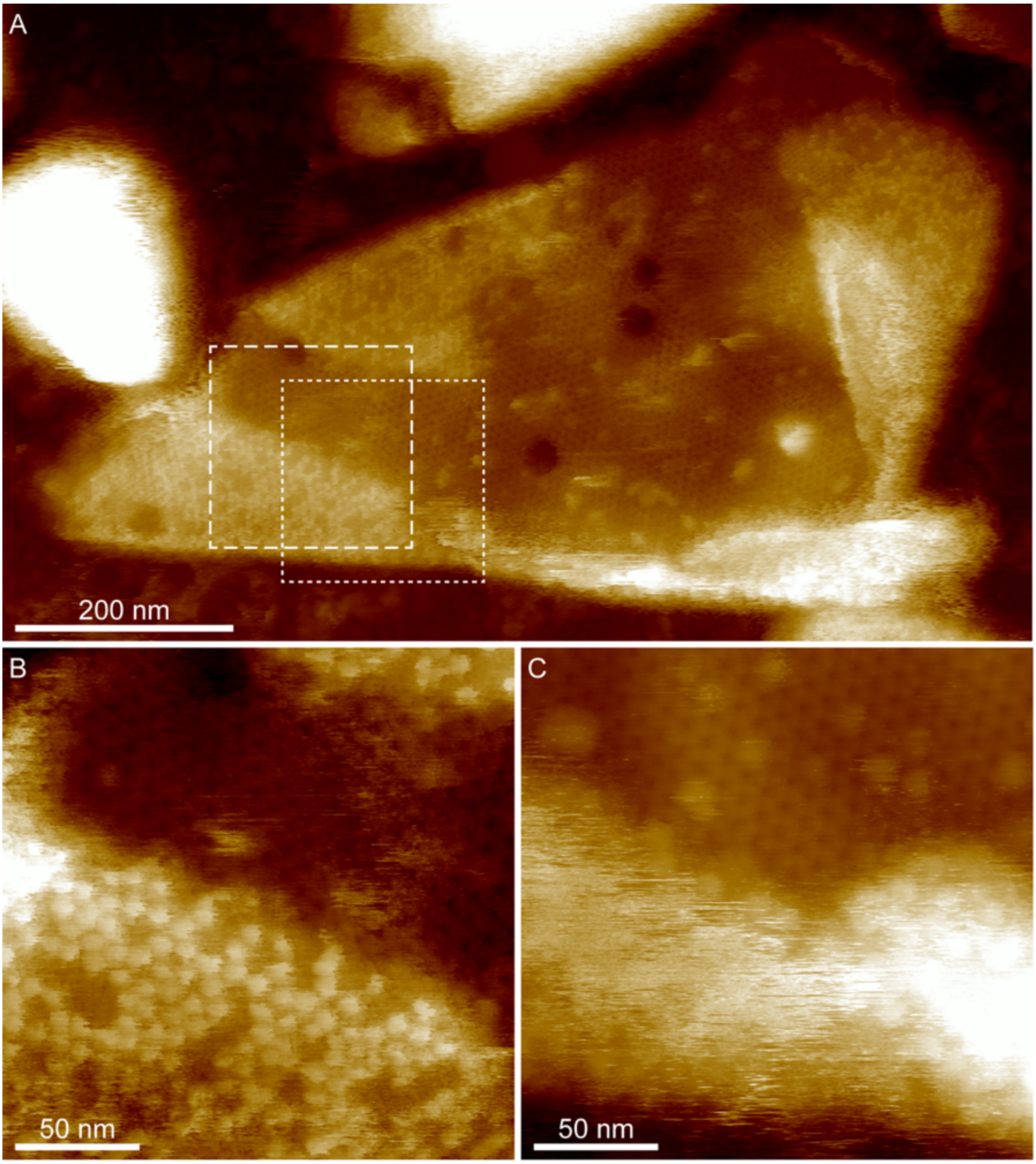


**Figure S4. AFM height images of hydrated exosporium reveal the underlying crystal lattice and a disordered surface.** (A) Exosporium fragment imaged with a free amplitude of ~2 nm and a relative setpoint of ~50% (high force). The honeycomb lattice is visible as a single layer area in the centre of the fragment. (B) Magnified image of the area shown by the dashed box in (A), also recorded under high force conditions. The honeycomb lattice is visible in the upper half of the image and a punctate lattice is visible on the folded over area in the lower half of the image. The apparent height difference between lower and upper half is 75-130 Å. (C) magnified area overlapping (B) and shown by dotted box in (A), imaged using low force (~1 nm free amplitude, ~90% setpoint). The honeycomb lattice remains visible in the upper half and the lower half, where the fragment has folded over, the opposing surface appears disordered. The height difference between lower and upper half is 200-450 Å.

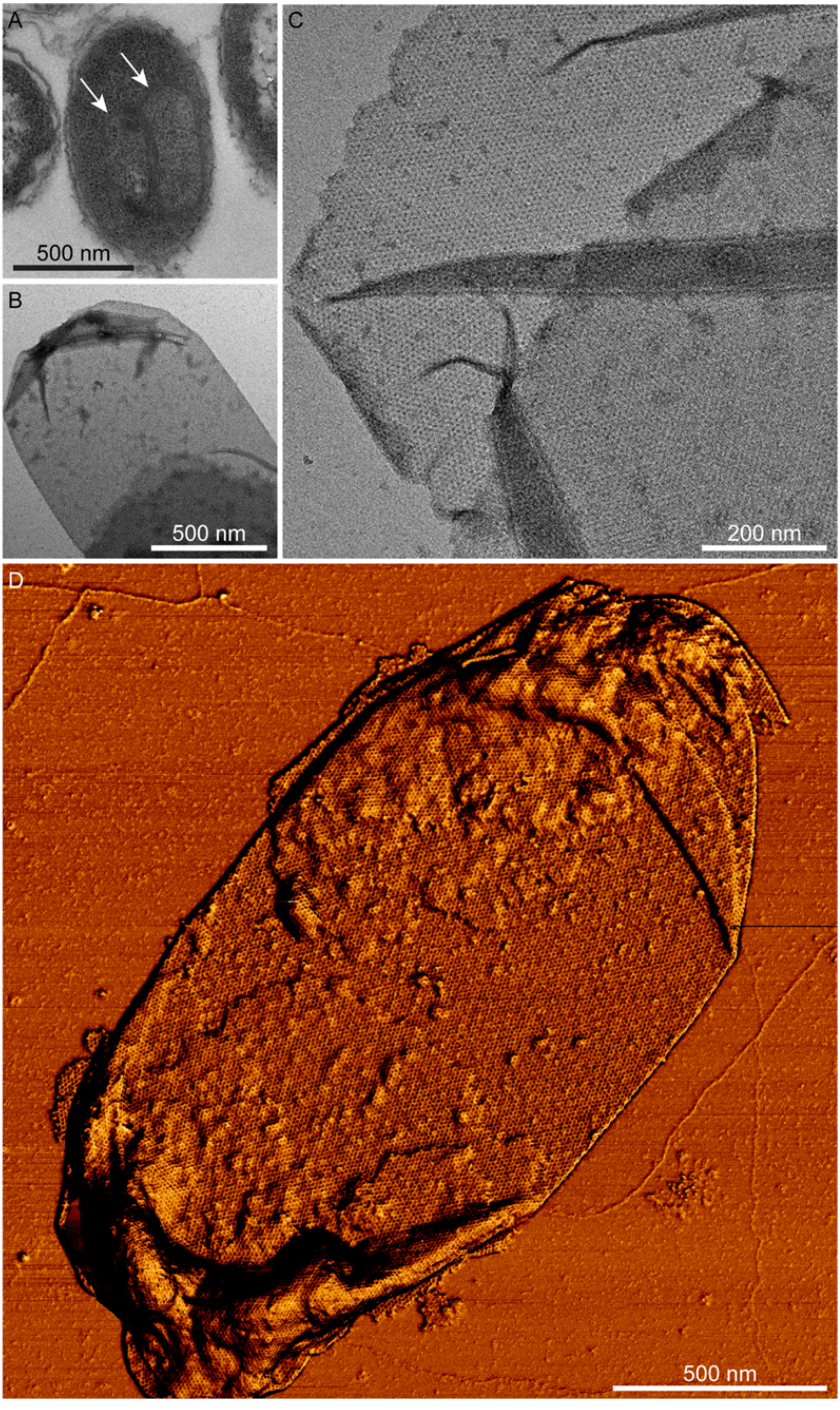


1

**(Previous page): Figure S5. Recombinant CsxA expressed in *E. coli* forms 2D crystals.** (A) A thin section electron micrograph showing formation of stacked layers within the cytoplasm (white arrows) of *E. coli* BL21 (DE3) pLysS cell expressing CsxA. (B) Electron micrograph of negatively stained sac-like 2D crystal of CsxA released from *E. coli* cell by sonication. (C) High magnification image of a broken sac, exposing a single 2D crystalline layer. (D) AFM phase image of purified intact sac shows the hexagonal arrays. Dark to bright variation in phase is 7.06°

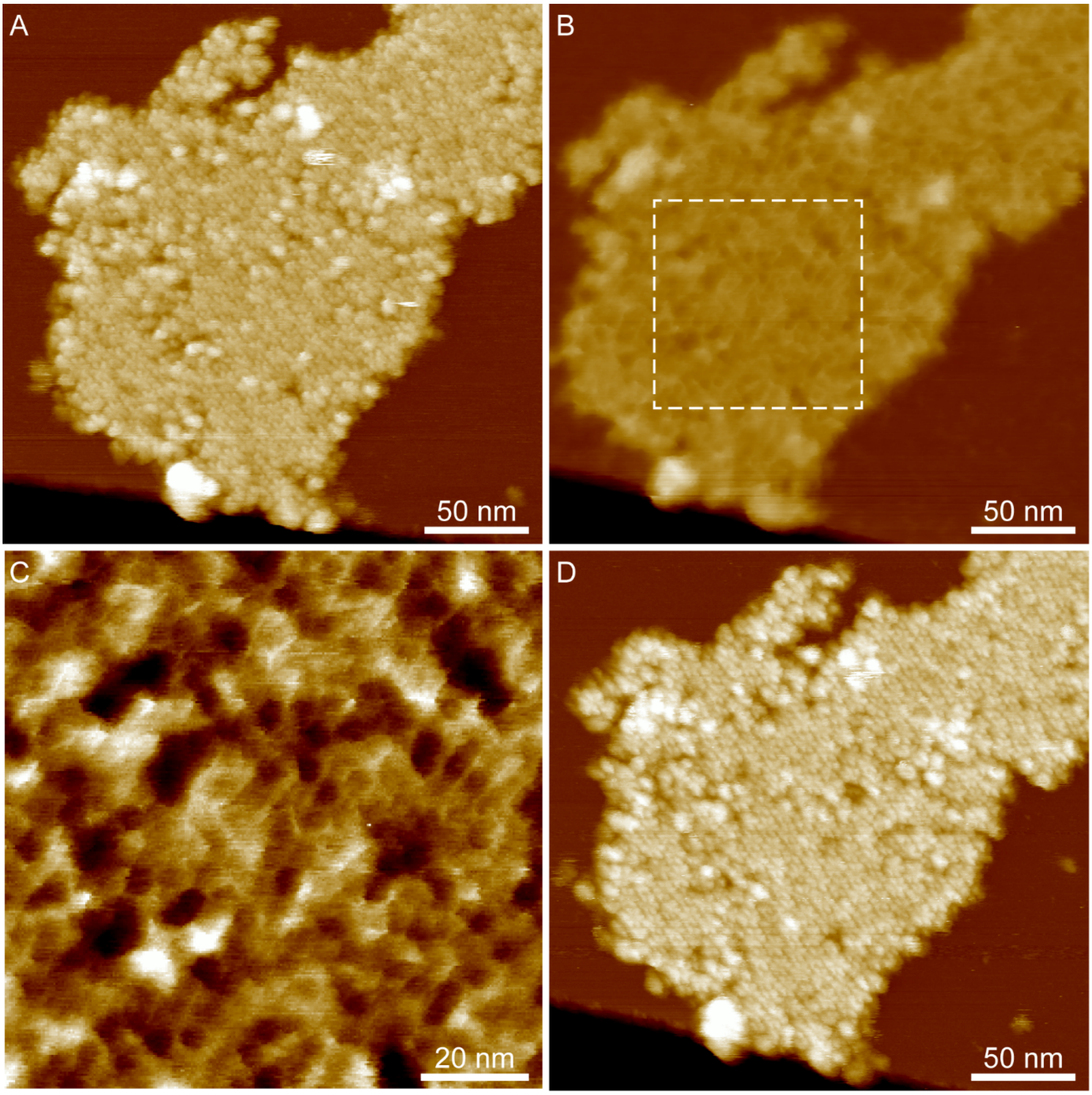


**Figure S6. AFM height images of CsxA showing a conformational change between wet and dry conditions.** (A) A recombinant CsxA 2D crystal imaged in water showing a hexagonal lattice of flower-like arrays with a lattice parameter of ~110 Å. Colour scale: 22 nm. (B) The same fragment of CsxA shown in (A) after drying, imaged in air. The flower like arrays are no longer visible and a hexagonal lattice of pores is present, with an apparent spacing of ~50 Å. Colour scale: 22 nm. (C) Magnified image of shown by the dashed box in (B). Colour scale: 2 nm. (D) The same fragment imaged in water after rehydration. The flower-like arrays become visible again, indicating that the conformational change is reversible. Colour scale: 22 nm.

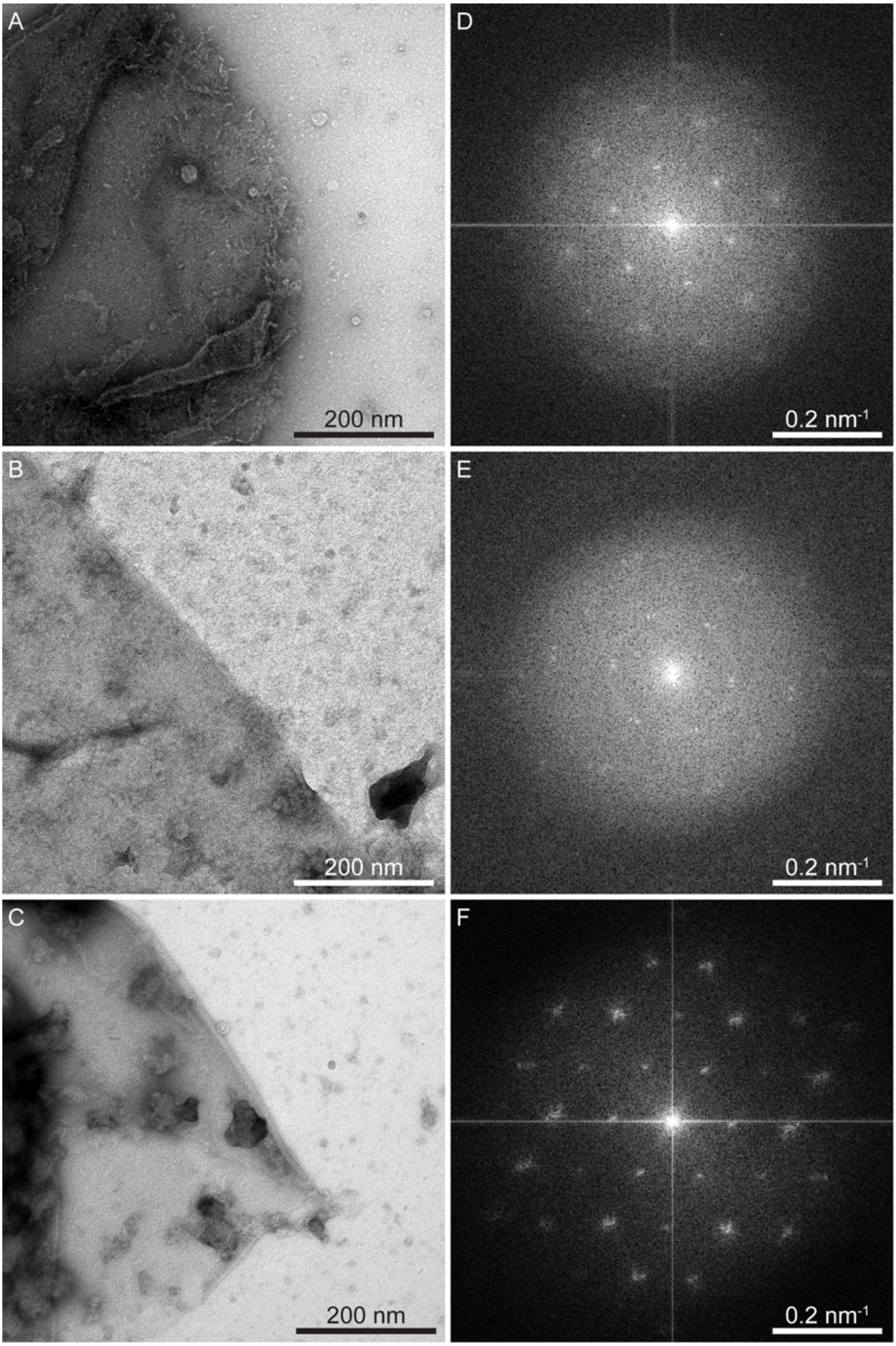


**Figure S7. CsxA crystals are thermally stable in non-reducing conditions.** (A)-(C) Recombinant CsxA crystals embedded in negative stain. (D)-(F) respective Fourier transforms from selected areas. (A), (D) Some crystals were found intact after incubation in 8M urea. (B), (E) Crystals were generally more disordered after incubation in 2M DTT at 25 °C. (C), (F) Crystals remained intact after incubation at 95°C for 60 mins.

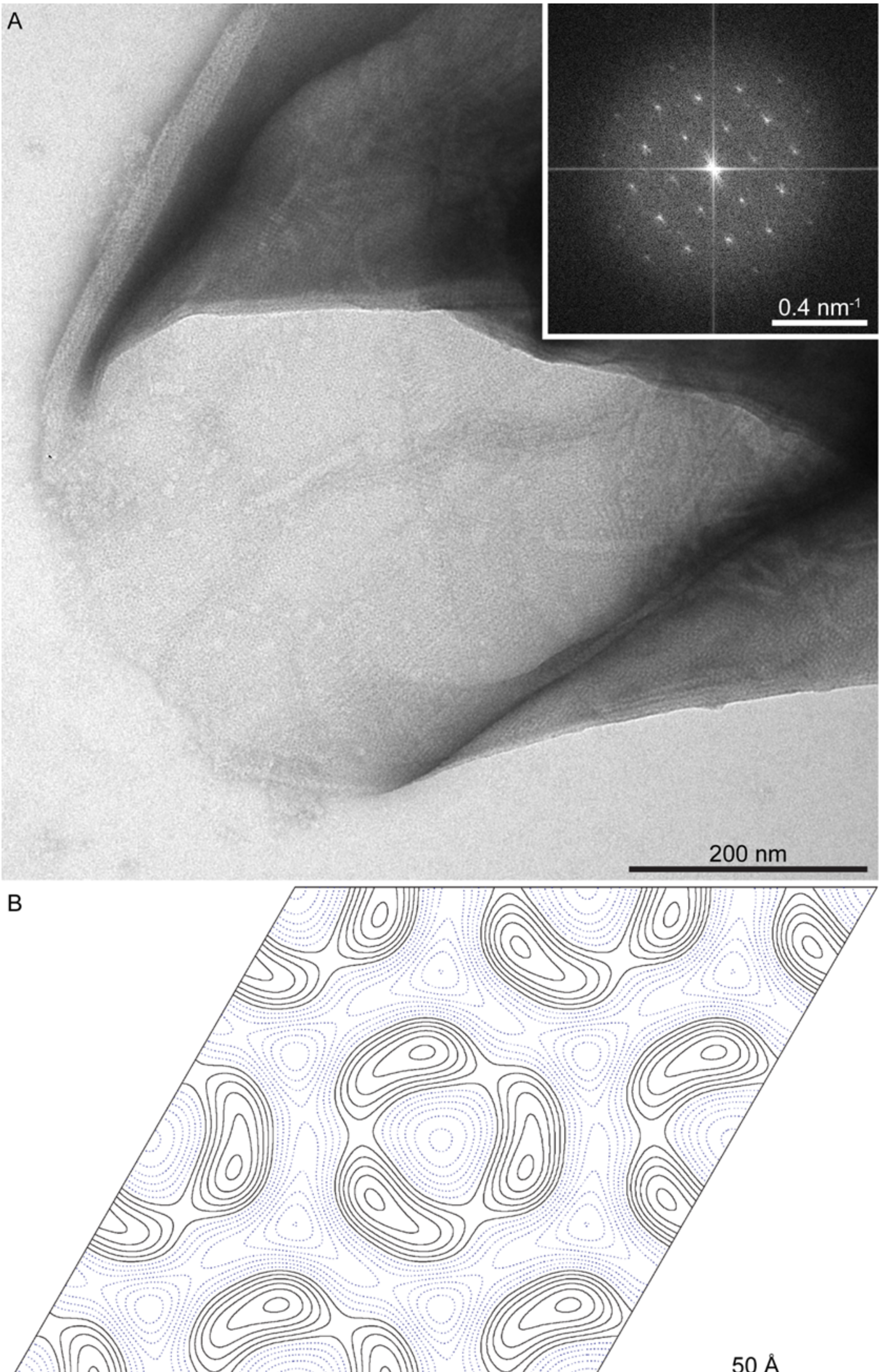


1

**(Previous page): Figure S8. 2D crystalline sheets from *csxA* spores display a different lattice from that of wild type exosporium.** (A) A high magnification image showing the lattice of a negatively stained crystal. Inset shows a computed diffraction pattern. (B) Projection map calculated with *p*3 symmetry imposed. Solid contours represent stain-excluding regions.

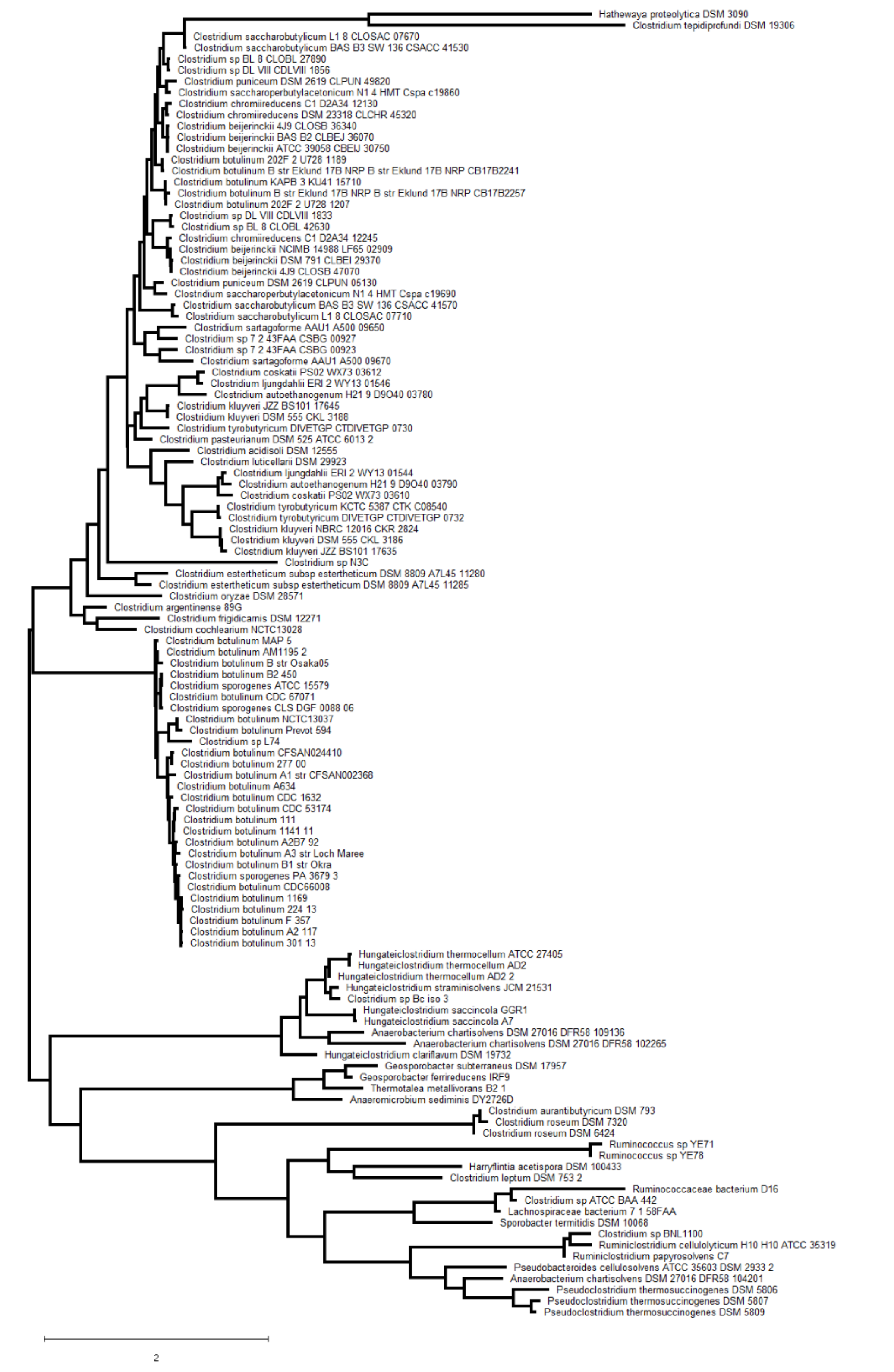


**Figure S9**. **CsxA homologues are found in other spore formers.** A maximum likelihood phylogenetic tree of CsxA homologues in spore formers constructed using RAxML (see supplementary methods). The scale bar indicates the number of amino acid substitutions per site represented by the indicated branch length.

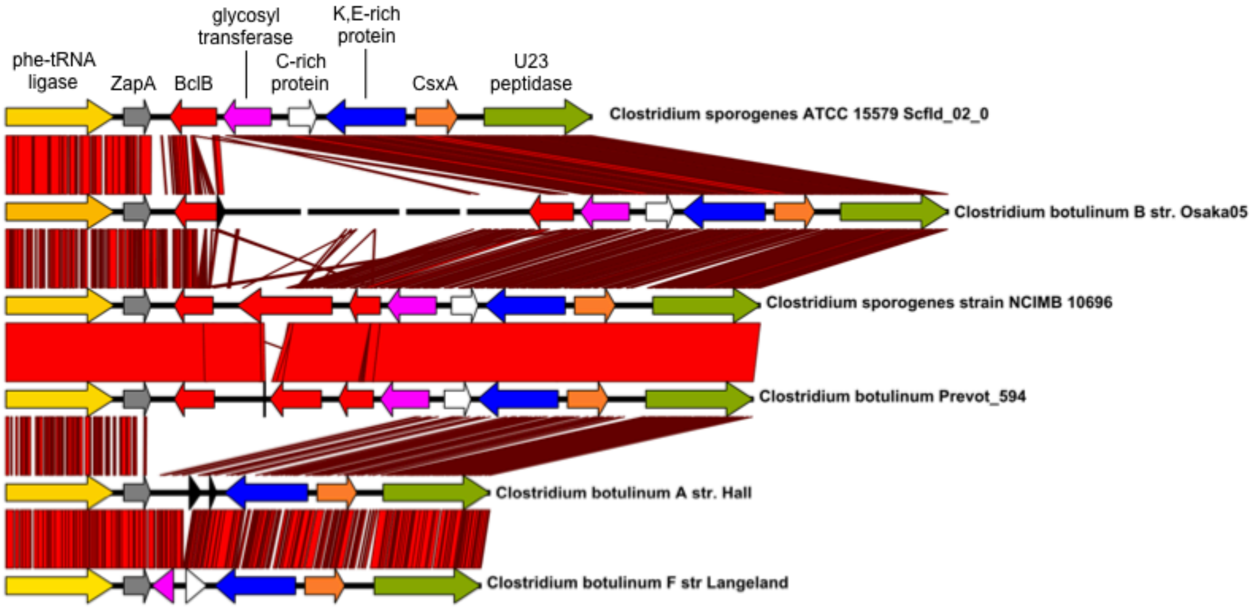


**Fig S10. Alignment of the genomic context of CsxA in *C. sporogenes* and *C. botulinum* Group I strains, from pheT to a U32 family peptidase gene.** Regions of sequence similarity between the strains are shown in red, with brighter shades indicating higher levels of sequence identity. Alignments were performed using NUCmer (1) and displayed using a custom version of the xBASE database (2). The region between zapA (grey: cell division gene) and the gene encoding a U32 peptidase (olive green) contains 7 ORFs in *C. sporogenes* NCIMB 10696, several of which correspond to protein sequences detected in the exosporium. The csxA gene is shown in orange, and is located at the right hand end of the cluster. Three ORFs in red at the left end each contain a central domain of collagen-like repeats (left to right these are *bclB* [CLSPO_c32280] and *bclA* [CLSPO_c32270] which encode proteins identified in the *C. sporogenes* exosporium (3), followed by CLSPO_c32260, which encodes a protein with a BclB-like C-terminal domain. The adjacent gene (shown in pink) encodes a glycosyl transferase, which may be responsible for glycosylation of one or more of the collagen-like repeat proteins, and is followed by an ORF encoding a cysteine-rich protein (white) and a highly conserved ORF (blue), organised divergently to csxA. *C. sporogenes* strain ATCC 15579 and *C.*

*botulinum* Osaka 05 have a smaller complement of *bcl*-like ORFs, and *C. botulinum* strains Hall and Langeland retain intact homologues of only *csxA* and its divergent partner.

1. Delcher AL, Salzberg SL, Phillippy AM (2003) Using MUMmer to identify similar regions in large sequence sets. *Curr Protoc Bioinformatics* 00:10.3.1–10.3.18.

2. Chaudhuri RR, et al. (2008) xBASE2: a comprehensive resource for comparative bacterial genomics. *Nucleic Acids Research* 36 (Database issue):D543–6.

3. Janganan TK, et al. (2016) Characterization of the spore surface and exosporium proteins of *Clostridium sporogenes*; implications for *Clostridium botulinum* group I strains. *Food Microbiol* 59:205–212.

### ***Supplementary methods***

*Spore preparation*

Spores of *C. sporogenes* NCIMB 701792 (NCDO1792) were grown on BHIS agar (Brain Heart Infusion agar supplemented with 0.1% L-cysteine and 5 mg/ml yeast extract) as described in (1, 2). Spores of *C. acetobutylicum* NCIB 8052, *C. tyrobutyricum* NCDO 1756, *C. puniceum* BL 70/20 and *C. pasteurianum* NCDO 1845 were grown on potato extract agar plates. Aliquots (100 µl) of an overnight cooked meat culture were spread on duplicate potato extract agar plates and incubated in an anaerobic chamber for three days at 30°C. Colonies were harvested from the surface of the plates using a sterile spreader and 1 ml of chilled saline. Plates were washed with a further 5 ml of saline and the spore suspension and washings from duplicate plates were combined into a centrifuge tube. Cultures containing spores, cells and cell debris were pelleted by centrifugation at 3000 x g for 15 min at 4°C. The pellets were washed three times in 10 ml saline with centrifugation at 2000 × g for 10 min at 4°C. Spores were separated from vegetative cells and debris on the basis of density. Washed pellets were resuspended in 1 ml saline, layered onto 10 ml 50% solution of sodium/meglumine diatrizoate (50 ml Urografin 370 (76%) mixed with 26 ml saline) and centrifuged at 5500 × g for 60 min at 4°C. The top layers were discarded, while the pellet was resuspended in 20 ml saline. Pellets were then washed a further four times in saline (2000 x g, 15 min, 4 °C), resuspended in 0.5 ml saline and stored at 2°C until required.

*csxA mutant construction*

Mutants of *C. sporogenes* strain ATCC15579 were generated using the Clostron system as previously described in (3). *C. sporogenes* strain NCIMB 701792 (NCDO1792) was not used for mutational studies as this strain is erythromycin resistant and not amenable to Clostron mutation. Target sites were identified in the *csxA* gene (gene; CLOSPO_00498 (*csxA*), insert site 345/346s) using the Perutka method (4) and mutants were generated as described in (5). Re-targeted introns were synthesized and ligated into the pMTL007C-E2 vector by DNA 2.0 (Menlo Park, USA). Plasmids were transformed into *E. coli* CA434 and then transferred by conjugation into *C. sporogenes* strain ATCC15579.

*Expression of csxA in E. coli*

Genomic DNA from *C. sporogenes* NCIMB 701792 was used as a template to amplify the *csxA* sequence by PCR using Phusion DNA polymerase (NEB), with forward primer ATCTA**CATATG**GCTATTAATTCAAAAGATTTTATTCCAC and reverse primer ATATT**CTCGAG**ATTATTAGTTATTACACTGCTAGTTATC. Oligonucleotides were designed based on the genome sequence of *C. sporogenes* PA 3679 strain. The amplified product was cloned between NdeI and XhoI sites in pET21a. The resulting plasmid was transformed into *E. coli* BL21 (DE3) pLysS for protein production.

For protein overexpression, *E. coli* cells were grown to optical density $OD_{600nm}$ of 0.5 in LB broth and induced with 1 mM IPTG for 3 hours at 37°C. Cells were harvested and resuspended in spore resuspension buffer (SRB) 25 mM Tris pH-8, 150 mM NaCl, and 1 mM PMSF and sonicated for 30 sec at an amplitude of 10 microns; sonication was repeated 3 to 5 times with 1 min intervals. The cell lysates were mixed with Ni-NTA agarose and incubated on an end over rotator at room temperature for 60 min. The Ni-NTA agarose was allowed to settle by gravity and the supernatant was discarded. The Ni-NTA beads were resuspended in 25 mM Tris pH-8, 150 mM NaCl buffer, packed in a gravity flow column and washed several times with buffer. The Ni-NTA agarose was transferred to a falcon tube and protein was eluted with 25 mM Tris pH-8, 150 mM NaCl and 1 M imidazole buffer. The eluted protein was centrifuged at 100,000 x g for 1 hour and the pellet containing CsxA 2D crystals was washed and resuspended in Tris buffer.

*Negative stain electron microscopy (EM)*

2 µl of spore suspension or sonicated *E. coli* cells were applied to glow discharged carbon coated copper palladium grids, incubated for 1 min, then washed, stained (20 s) with uranyl formate (0.75%) and vacuum dried. For CsxA crystals extracted from sonicated *E. coli* cells, after sample incubation, the grids were washed once in distilled water before being stained. Grids were examined in a Philips CM100 operating at 100 kV. Micrographs were recorded under low dose conditions on a Gatan MultiScan 794 1k x 1k CCD camera at nominal 52,000x magnification with 0.5-1.2 µm underfocus and at specimen tilts over a range of ± 55° in 10° steps.

*Electron cryomicroscopy (CryoEM)*

Recombinant CsxA 2D crystals purified as above were resuspended in Tris buffer (pH 7.5, 150 mM NaCl and 1 mM EDTA) and loaded onto glow discharged carbon-coated molybdenum grids or Quantifoil R2/2 grids with carbon support and incubated for 1 min. Grids were blotted and plunge frozen into liquid ethane using an FEI Vitrobot plunge freezer or Leica plunge freezer; blotting times used were 25s and 4-6s respectively.

Samples were examined on a Philips CM200 FEG EM, equipped with an Oxford Instruments CT3500 cold stage, or a FEI Tecnai Arctica FEG EM both operated in low-dose at 200kV and liquid nitrogen temperature. Data collected on the CM200 were at a nominal magnification of 66,000x and recorded on a CCD Gatan UltraScan 890 (US4000SP) 2k x 2k camera, with defocus values of ~1 µm and 0.5 s exposure. On the Tecnai Arctica micrographs were recorded at magnifications of 39,000-78,000x on a Falcon 3 direct electron 4k x 4k detector at defocus values of ~1.5-6.5 µm with either a 1 s single exposure or 2 s exposure over 79 frames and motion corrected with MotionCorr2.

*Electron microscopy of E. coli cell sections*

*E. coli* cells overexpressing CsxA were pelleted (100 µl) and fixed with fresh 3% glutaradehyde in 0.1 M phosphate buffer overnight at 4°C. The sample was then washed in 0.1 M phosphate buffer two times at 30 min intervals at 4°C. Secondary fixation was carried out in 2% aqueous osmium tetroxide for 2 h at room temperature and the sample washed as above. The sample was dehydrated using a series of ethanol washes and finally dried over anhydrous copper sulphate for 15 min. The sample was then placed in two changes of propylene oxide for 15 min. Infiltration was achieved by placing the sample in a 50/50 mixture of propylene oxide/araldite resin overnight at room temperature. The sample was left in full strength araldite resin for 6-8 hours at room temperature after which it was embedded in fresh araldite resin for 48-72 hours at 60°C. Ultrathin sections, approximately 70-90 nm thick, were cut on a Reichert Ultracut E ultramicrotome and stained for 25 min with 3% uranyl acetate followed by staining with Reynold's lead citrate for 5 min. Sections were examined on a FEI Tecnai 120 G2 Biotwin electron microscope at 80 kV with a Gatan Orius SC 1000B digital camera (bottom mounted).

*Image processing*

Electron micrographs were processed within the *2dx* software suite (6) based on the MRC suite of programs (7). Images were subjected to two cycles of unbending. For all subsequent analysis *p*6 symmetry was enforced. Phase origins for individual images were refined against each other using ORIGTILTD, sequentially adding images of increasing tilt angle to the refinement. Initial estimates of tilt angle were made from more highly tilted members of a series using EMTILT (8). The common phase origin was found by comparing the phases of the reflections on each image within a z* value of 0.01 $Å^{-1}$ to those of all the other images. At least two cycles of refinement of the phase origin, tilt angle and tilt axis for each image, were performed. The program LATLINE (9) was used to determine interpolated amplitudes and phases on a regular lattice of 1/270 $Å^{-1}$ in z* for data up to 1/25 $Å^{-1}$ resolution. A generous real-space envelope of approximately twice the estimated exosporium thickness (70 Å) was applied as a constraint. The output interpolated lattice lines were used as references for two cycles of crystal tilt and phase origin refinement. The variation of amplitude and phase along 0,0,l was estimated from the maximum contrast on each Z-section in the 3D map (10). Density maps were calculated within the CCP4 suite of crystallography programs (11). 3D surface representations were rendered with CHIMERA (12).

For frozen hydrated CsxA crystals we merged and averaged 14 images in *2dx* (6) to 9 Å resolution. A negative temperature factor (B-factor) was estimated and applied to the projection map by scaling averaged image amplitudes against bacteriorhodopsin electron diffraction amplitudes (13).

*Atomic Force Microscopy (AFM)*

Exosporium fragments were prepared as described in (1). 10 µl of spore fragment solution (concentration 2.2 mg/ml) was diluted in 100 µl citric acid/sodium phosphate buffer, 150 mM KCl (pH 4) and incubated on poly-d-lysine coated coverslides (Corning BioCoat) for 30 min at room temperature.

2D crystals of CsxA were prepared for AFM by incubating 2-5 µl of crystals in storage buffer (20 mM Tris, 150 mM NaCl, (pH 8)) on freshly cleaved mica for ~30 min at room temperature. After binding, all samples were washed with 10x 1 ml HPLC grade de-ionized water and either imaged in water or dried with filtered nitrogen and imaged in air.

Imaging in air was performed using a JPK NanoWizard Ultra AFM in AC mode with TESPA V2 cantilevers (nominal stiffness 37 N/m, nominal resonant frequency 320 kHz) in a home built vibration and acoustic isolation system. The free amplitude was approximately 8 nm and the relative setpoint 90-95%. Imaging in water was performed using a Bruker Dimension FastScan AFM in Tapping Mode with FastScan D cantilevers (nominal stiffness 0.25 N/m, nominal resonant frequency (in water) 100 kHz). The cantilever holder was washed in household detergent, isopropanol and pure water before each experiment. Imaging was performed in a small water drop (~200 μl). The free amplitude was approximately 1 nm and the relative setpoint 80-90%.

To minimize the effects of thermal drift, the sample was placed into the AFM and allowed to settle for approximately 1 h (experiments in water) and 2-24 h (experiments in air) with the isolation hood closed. In all cases, the probe was tuned close to the surface after engagement to compensate for squeeze film damping. The feedback gains, scan rate, pixel density and amplitude setpoint were adjusted for optimal image quality while scanning. To ensure accuracy of all measurements, scanner calibration was checked using a 3 μm pitch and 180 nm depth grating and found to be within 1% for all 3 axes.

Images were processed (flattening, planefitting) and analyzed using JPK DP software, Gwyydion and NanoScope Analysis.

*Phylogenetic Analysis of CsxA homologues*

To identify homologues of *csxA*, all available complete and draft genome sequences of spore-forming bacteria were downloaded from GenBank, and the annotated protein sequences were extracted and converted into a BLAST database for each strain. These were interrogated with BLASTp, using the CsxA sequence of *Clostridium sporogenes* ATCC 15579 as a query. Any hits with >20% amino acid identity to CsxA over at least 50% of the length of the query sequence were retained, resulting in the identification of 265 potential CsxA homologues. Duplicate sequences were removed, giving a dataset of 117 distinct protein sequences of CsxA homologues. These were aligned using Muscle version 3.8.31 (14), and a phylogenetic tree was constructed with RAxML version 8.2.12 (15) using the VT model of amino acid substitution (16), which was selected as the best scoring amino acid model by RAxML, and a gamma model of rate heterogeneity.


1. Janganan TK*, et al.* (2016) Characterization of the spore surface and exosporium proteins of *Clostridium sporogenes*; implications for *Clostridium botulinum* group I strains. *Food microbiology* 59:205-212.
2. Smith CJ, Markowitz SM, & Macrina FL (1981) Transferable tetracycline resistance in *Clostridium difficile*. *Antimicrobial agents and chemotherapy* 19:997-1003.
3. Brunt J*, et al.* (2014) Functional characterisation of germinant receptors in *Clostridium botulinum* and *Clostridium sporogenes* presents novel insights into spore germination systems. *PLoS pathogens* 10:e1004382.
4. Perutka J, Wang W, Goerlitz D, & Lambowitz AM (2004) Use of computer-designed group II introns to disrupt *Escherichia coli* DExH/D-box protein and DNA helicase genes. *Journal of molecular biology* 336:421-439.
5. Heap JT*, et al.* (2014) Spores of Clostridium engineered for clinical efficacy and safety cause regression and cure of tumors in vivo. *Oncotarget* 5:1761-1769.
6. Gipson B, Zeng X, Zhang ZY, & Stahlberg H (2007) 2dx--user-friendly image processing for 2D crystals. *Journal of structural biology* 157:64-72.
7. Crowther RA, Henderson R, & Smith JM (1996) MRC image processing programs. *Journal of structural biology* 116:9-16.
8. Shaw PJ & Hills GJ (1981) Tilted Specimen in the Electron-Microscope - a Simple Specimen Holder and the Calculation of Tilt Angles for Crystalline Specimens.

*Micron* 12:279-282.

***Supplementary tables***

**Table S1. 3D merging statistics for native exosporium reconstruction in negative stain**

| | |
|---|---|
| Resolution limit | 25 Å |
| Number of structure factors | 1086 |
| Overall R-factor | 0.33 |
| Overall phase residual | 27.2° |

**Table S2. The internal phase residuals determined after the imposition of all possible two-sided plane groups, calculated from one of the micrographs of frozen hydrated CsxA crystals**

Internal phase residuals were determined from spots of IQ1 to IQ5 to 7 Å resolution. The values marked * are good candidates for the symmetry as the experimental phase residual is close to or better than that expected based on the signal-to-noise ratio.

| Two-sided plane group | Phase residual versus other spots (90° random) | Number of comparisons | Target residual based on statistics taking Friedel weight into account |
|---|---|---|---|
| *p*2 | 31.8* | 57 | 37.5 |
| *p*3 | 13.9* | 82 | 25.7 |
| *p*312 | 49.1 | 173 | 26.3 |
| *p*321 | 40.0 | 178 | 26.6 |
| *p*6 | 17.5* | 221 | 28.7 |
| *p*622 | 46.2 | 408 | 27.3 |

**Table S3. 3D merging statistics for CsxA crystals in negative stain**

| | |
|---|---|
| Resolution limit | 25 Å |
| Number of structure factors | 962 |
| Overall R-factor | 0.33 |
| Overall phase residual | 36.4° |

**Table S4. Phase residuals in resolution shells for *p*6 averaged Fourier terms of frozen hydrated CsxA crystals**

| Resolution shell (Å) | Number of independent reflections | Mean value of phase error against symmetry imposed phase of 0° or 180° (45° is expected for random phases) | Standard error (°) |
|---|---|---|---|
| 200-15 | 25 | 11.0 | 4.0 |
| 15-12 | 14 | 31.9 | 6.9 |
| 12-10 | 13 | 25.3 | 4.4 |
| 10-9 | 18 | 25.9 | 5.9 |

**Table S5. Effect of incubation of CsxA crystals under different conditions.**

Crystalline order: high- diffraction beyond second order; low- diffraction up to first order. Crystal abundance: high- more than 5 crystals/grid square; low- fewer than 1 crystal/grid square

| | No treatment | 8M urea | 95°C, 60 mins. | 2M DTT, 25 °C | 2M DTT, 95°C |
|---|---|---|---|---|---|
| Crystals observed | yes | yes | yes | yes | no |
| Crystalline order | high | low-high | high | low | - |
| Crystal abundance | high | low | high | low | - |

**Supplementary data set 1. Cysteine-rich proteins are clustered with collagen-like repeat proteins across spore formers.** All available complete and draft genomes of spore-forming bacteria were interrogated for the presence of collagen-like proteins, identified by a HMMER3 search using the collagen triple helix repeat domain (Pfam accession PF01391.18). The table lists all cysteine-rich proteins (defined as $\log_{10}$(number of Cys residues/protein length) > -1.25, equivalent to a cysteine content of >5.6%) encoded by genes with a midpoint within 20kb of the midpoint of a gene encoding a collagen-like protein.
